# Decomposition of task-specific responses in the multiple demand network

**DOI:** 10.64898/2026.06.21.732474

**Authors:** Neda Afzalian, Mohammad Ebrahim Katebi, Reza Rajimehr

## Abstract

Neuroimaging evidence suggests that a distributed network of brain areas, known as the multiple demand (MD) network, is consistently recruited across a wide range of cognitive tasks. The MD network includes specific areas in prefrontal and parietal cortices. However, the fine-grained organization of this network is poorly understood. Here we aim to comprehensively characterize the functional subdivisions within the MD network using a naturalistic fMRI paradigm. 20 subjects were instructed to perform 14 different tasks on a set of movie stimuli. These tasks were designed to target a wide range of cognitive domains including visual, spatial, categorical, emotional, auditory, linguistic, social, and semantic processes. fMRI data were also collected while subjects passively watched the movies. The MD network was first delineated by localizing cortical areas that were significantly more active in all 14 tasks compared to the passive-viewing condition. The principal component analysis was then applied on task-specific responses of cortical points within the MD network. The first component was correlated with activities for all tasks, and its spatial map revealed the core, highly multimodal subregions of the MD network. The other components showed preferences for a subset of tasks. In particular, the second component revealed a sharp distinction between MD regions that were preferentially active in visual/spatial versus linguistic/semantic tasks. A graph analysis on the entire cortex also showed a large-scale distinction between visual/spatial and linguistic/semantic areas, with MD regions linking the two communities of cortical areas. Overall, our results provide new insights into how the MD network and its fine-grained architecture contribute to the human intelligent behavior.

## Introduction

Human cognition depends on the rapid and flexible coordination of multiple processes to meet dynamic goals and environmental demands [1–3]. This flexibility arises from the brain’s capacity to integrate information across distributed cortical systems, linking perceptual and higher-order association areas to support fluid intelligence and adaptive behavior [4–6]. The MD network is central to this integrative capacity [7–9]. It is consistently activated across a broad spectrum of cognitively demanding tasks [10] and supports a range of distinct executive functions, including cognitive control [11], cognitive integration [12], working memory [13,14], selective attention [15,16], and inhibition [17, 18]. Within large-scale network taxonomies, the MD network closely aligns with the frontoparietal control network (FPN) [19–22], with substantial anatomical and functional convergence between the two [23]. Key MD regions include the dorsolateral prefrontal cortex, inferior frontal junction, anterior insula, and parietal territories surrounding the intraparietal sulcus and superior parietal lobule [24, 25].

The dominant account holds that the MD network operates as a homogeneous, domain-general system whose engagement is driven by cognitive control demand rather than task content or modality [9, 26–29]. Consistent with this view, lesions to any major MD region yield broad impairments in executive function and fluid intelligence, underscoring the system’s distributed yet functionally unified contribution to goal-directed behavior [30–35]. Neuroimaging studies further show compensatory engagement of intact MD regions following focal damage, highlighting the cooperative organization of the system [36, 37]. Resting-state functional connectivity analyses demonstrate dense coupling among MD regions, consistent with a tightly integrated control network [5, 23, 25]. Task-based multivoxel-pattern analyses show that MD regions flexibly encode whatever is currently task-relevant, with decodable information spanning tactile, visual, auditory, rule-based, and motor domains [38–45]. Large multi-task analyses show that MD regions reorganize their interactions with other brain systems according to task demands while consistently playing a central coordinating role across tasks [46]. Together, these findings characterize the MD network as a unified, flexible architecture that supports domain-general executive control across diverse cognitive demands.

Although the MD network is often characterized as a unitary, domain-general control system, converging evidence suggests a more differentiated internal organization. Meta-analytic studies report partially dissociable recruitment patterns for working memory, inhibition, and attentional shifting within frontoparietal MD territories [17, 47, 48]. Related distinctions have been observed in inferior parietal cortex, where neighboring subregions show preferential engagement for attentional, semantic, or social-cognitive demands [49]. Complementing these findings, recent high-resolution analyses indicate that updating, shifting, and inhibition elicit overlapping yet reliably distinguishable activation profiles across MD territories and their adjacent networks [50]. Evidence for such heterogeneity has also emerged from resting-state studies, particularly those emphasizing dominant network components rather than coarse parcellations [20], as well as studies explicitly accounting for inter-individual variability, which can obscure fine-grained structure in group-averaged analyses [51, 52]. Within this literature, analyses focusing on frontoparietal subnetworks suggest that differentiation becomes apparent when MD regions are considered in relation to other large-scale systems (e.g., default mode and dorsal attention networks) [53, 54]. Other connectivity-based accounts, however, emphasize a tightly integrated MD core, proposing that functional diversity arises primarily within an extended MD system surrounding this core rather than within the core itself [23, 55].

To resolve the conflicting evidence concerning both functionally uniform and differentiated accounts of the MD network, it is essential to examine its fine-grained functional organization using a sufficiently rich and systematically varied set of tasks. Limitations of previous work, including restricted task diversity, heterogeneity in sensory inputs, and insufficient spatial resolution, have constrained the field’s ability to adjudicate between these competing views. A more comprehensive and fine-grained characterization of MD activity across diverse cognitive demands is therefore needed to determine whether the network contains meaningful internal specialization or primarily operates as a domain-general control system.

We aim to address these challenges by employing a naturalistic fMRI paradigm in which subjects view identical movie clips while performing tasks designed to engage perceptual, affective, social, and semantic processes. Holding the sensory input constant minimizes stimulus confounds and allows us to examine how the MD network is engaged in both domain-general and task-specific contexts. We first localize the MD network using this rich dataset, providing an alternative approach that complements traditional task-based localizers. Building on this localization, we then investigate whether the MD network exhibits functional subdivisions that are selectively engaged by different cognitive demands and further explore how these subdivisions are embedded within large-scale cortical networks. Finally, we examine how the organization of the MD network relates to individual differences in cognitive performance, offering insight into how its structure supports diverse cognitive functions across domains.

## Results

To investigate the fine-grained organization of the MD network, we conducted an fMRI experiment with 15 conditions: 14 task conditions targeting perceptual, emotional, social, and semantic domains, and one passive-viewing condition (Figure 1A,B). Across all conditions, subjects viewed the same set of naturalistic movie clips, carefully constructed to incorporate content from all fourteen cognitive domains while maintaining constant sensory input (see Methods for details). After each movie clip, subjects completed a brief two-choice question to verify task-specific engagement (Figure 1C, Supplementary Figure S1). Overall, participants performed reliably well across the tasks, with the exception of the scene condition, which showed lower accuracy (Figure 1D). In the passive-viewing condition, subjects were cued to watch the clips, and at the end of the clip, respond to a simple arithmetic question which was unrelated to the movies.

**Figure 1:**
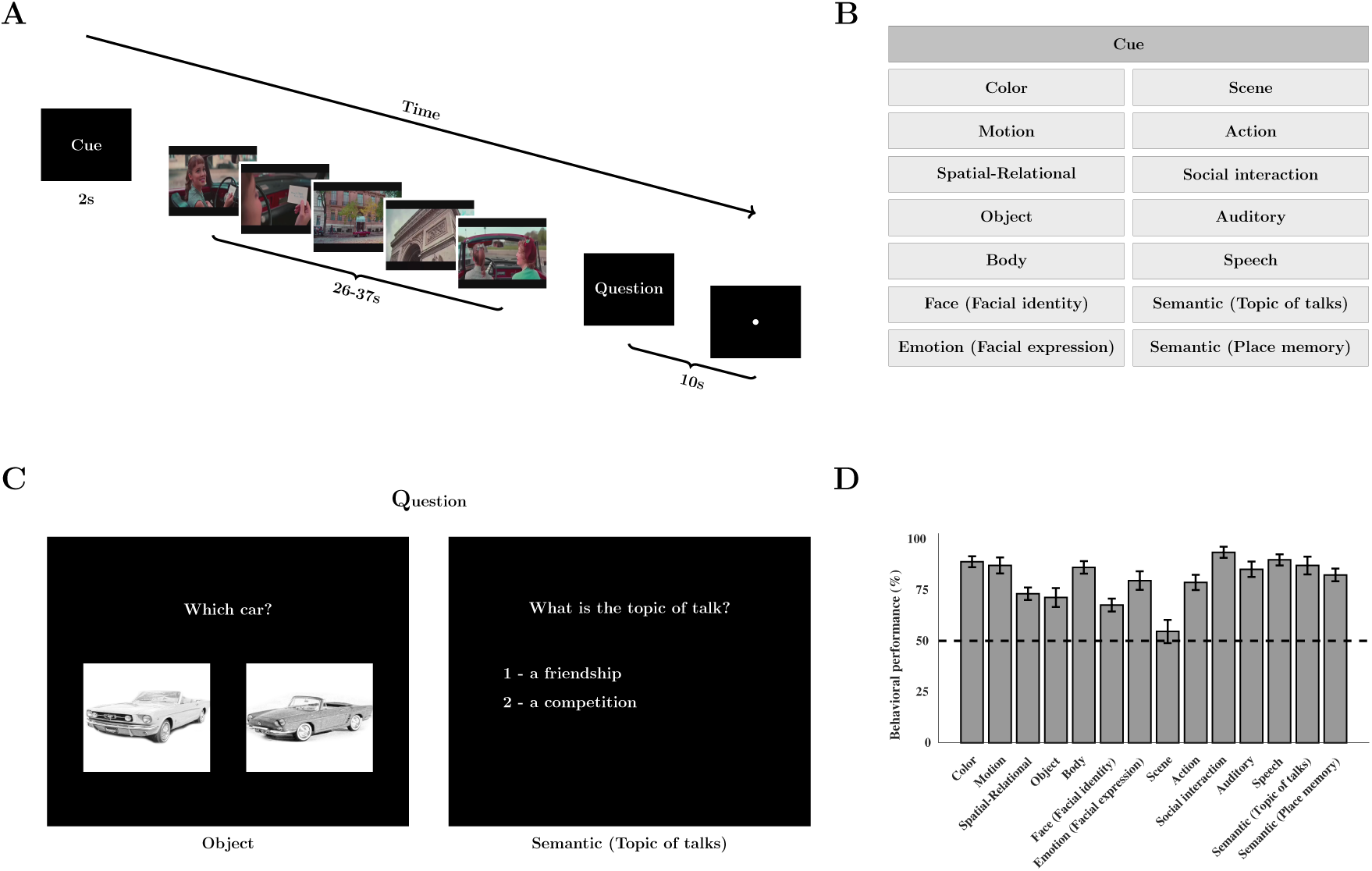
Experimental paradigm and behavioral performance. **(A)** A schematic representation of the events within a single task-specific trial. Each trial began with a cue event (2 s), consisting of a 1-second cue indicating the target cognitive domain (e.g., face, object, speech), followed by a 1-second blank screen. The cue event was immediately followed by a movie event (26-37 s) in which subjects focused their attention on the cued domain within the movie clip. At the end of the clip, a question event (up to 10 s) prompted subjects to answer a two-choice question related to the attended domain by pressing a key on a response box. Blank trials of varying durations were interspersed throughout the run. **(B)** Cognitive domains and their associated cues. The table lists the 14 cognitive domains targeted by the domain-specific tasks. **(C)** Examples of the two types of question events: left, a two-choice question with pictorial answers (used for domains such as face and object); right, a two-choice question with text-based answers (used for domains such as speech and semantic content). **(D)** Task performance across cognitive domains. The bar plot shows the average accuracy (± standard error of mean) of subjects for the two-choice question in each of the 14 cognitive domains. Dashed line indicates chance-level performance (50%).

We then estimated task-specific neural responses at the subject level. fMRI data were first preprocessed and then modeled using a general linear model (GLM) for each of the 15 conditions. For each run, activation maps were obtained by contrasting the movie-viewing periods versus the blank intervals, capturing task-specific responses relative to the non-stimulus blank. The resulting maps were subsequently aligned across subjects using multimodal surface-based registration informed by cortical folding, myelin content, and functional connectivity. These subject-level activation estimates formed the basis for the group-level analyses aimed at localizing the MD network and characterizing its functional organization across cognitive domains.

### Localization of the MD Network

The MD network was identified using a group-level analysis comparing activation across all cognitive tasks versus the passive-viewing condition. For each cortical vertex, we calculated the probability of tasks showing significantly higher activation in the task versus the passive-viewing condition (paired-sample t-test, *p <* 0.05). This procedure identified regions consistently engaged across diverse tasks and produced a robust probabilistic representation of the MD network (Figure 2A). The resulting map revealed multiple distinct activation patches distributed across the cortical surface. The most prominent patches were observed in the parietal and frontal cortices, corresponding to known anatomical locations of the MD network [23]. In the parietal cortex, three patches were identified: one on the lateral surface and two on the medial surface. The lateral parietal cortex (LPC) patch was located within the intraparietal sulcus. The medial parietal patches were located near the parieto-occipital sulcus (POS) and the retrosplenial cortex (RSC). The frontal cortex also contained two prominent patches, located within the inferior frontal cortex and the frontal pole. Additional, less prominent patches were observed in the anterior insula, dorsomedial frontal cortex, and lateral temporal cortex.

**Figure 2:**
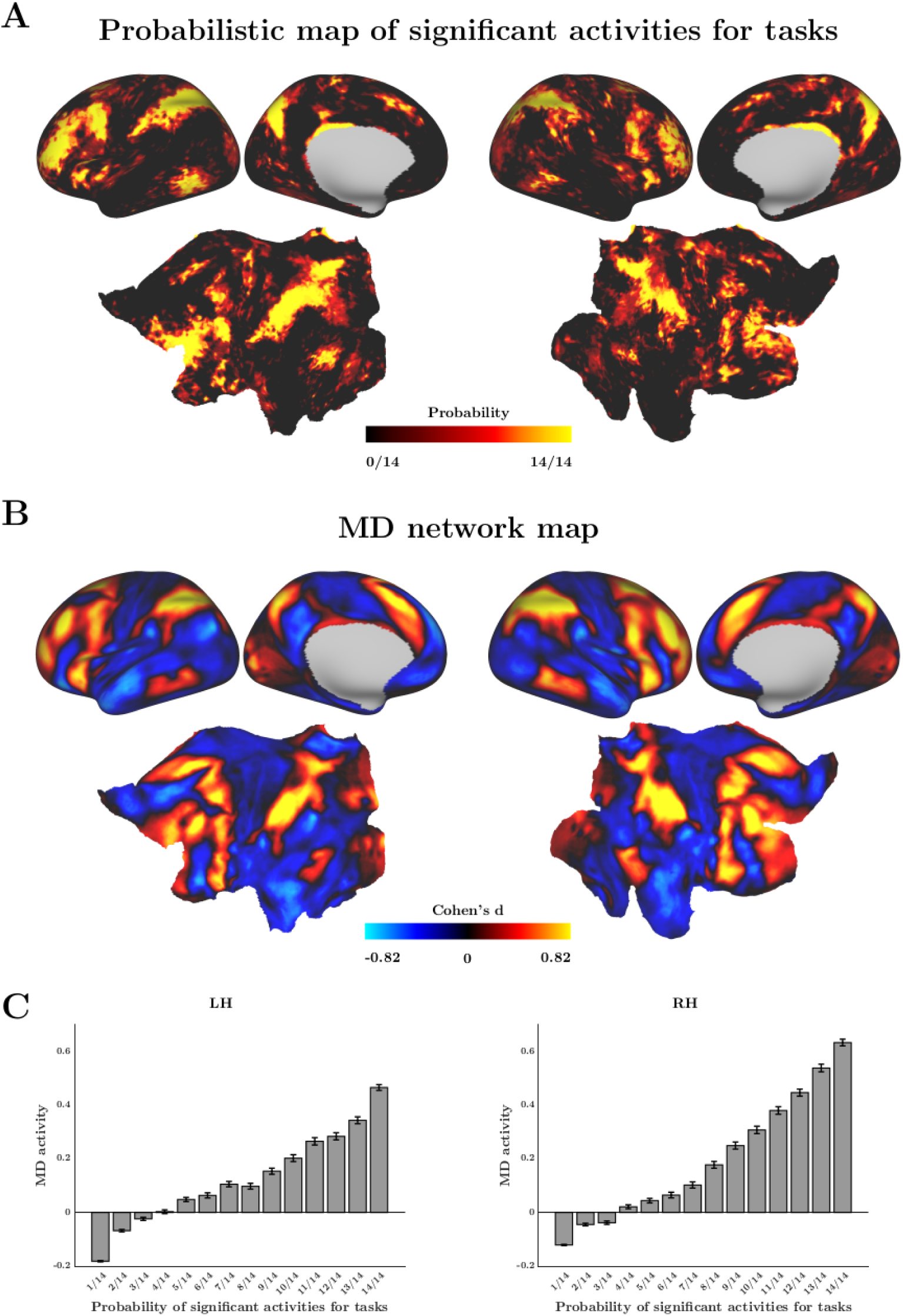
Localization of the MD network using 14 movie-based tasks. **(A)** Probabilistic map of task-evoked activations. This map shows the probability that each cortical vertex was significantly active across all 14 task conditions when contrasted with the passive-viewing condition. The colorbar indicates the proportion of tasks with significant activation, ranging from black (0/14) to yellow (14/14). **(B)** HCP-based MD map. This map shows group-average Cohen’s d values from three task contrasts within the HCP dataset: 2-back vs. 0-back working memory, hard vs. easy relational reasoning, and math vs. story in the language task. In panels A and B, left and right hemisphere maps are shown on the left and right, respectively. The top row shows lateral and medial views of an inflated cortical surface, while the bottom row shows flat maps. **(C)** The barplots show the average MD activations (Cohen’s d values) from the HCP dataset, with data grouped by the probability of activation determined in panel A. Error bars indicate standard error of the mean across vertices.

To validate this localization, we replicated the previously conducted analysis by Assem et al. [23] using large-scale Human Connectome Project (HCP) data from 1,080 subjects. Three well-established task contrasts were used to define the MD network: 2-back vs. 0-back working memory, hard vs. easy relational reasoning, and math vs. story in the language task. The resulting group-average MD map, obtained by averaging activities across these contrasts, closely matched the spatial organization observed in our probabilistic map (Figure 2B).

Building on this cross-dataset validation, we quantified vertex-wise correspondence between the probabilistic and HCP-based MD maps (Figure 2C). Vertices in our map were stratified according to their probability of task engagement, and for each probability tier, the mean activation value was extracted from the HCP-based map. This analysis revealed a systematic relationship: as the probability of task-related engagement increased, average HCP activation enhanced accordingly. This convergence across independent datasets demonstrates the robustness and reproducibility of our MD network localization. The effect was evident in both hemispheres, with a more pronounced gradient in the right hemisphere.

We then refined the distributed probabilistic map obtained from the vertex-wise analysis to delineate the core regions of the MD network. Although the initial probabilistic map showed substantial overlap with well-established MD network regions, it also included vertices distributed across additional cortical regions. To focus subsequent analyses on the most stable and functionally cohesive components of the network, we repeated the analysis at the parcellation level using parcels derived from the HCP multimodal parcellation (MMP) [56]. Parcels that were consistently activated across all 14 tasks in the task vs. passive-viewing comparison (paired-sample t-test, *p <* 0.05) were retained for subsequent analyses (Figure 3A).

**Figure 3:**
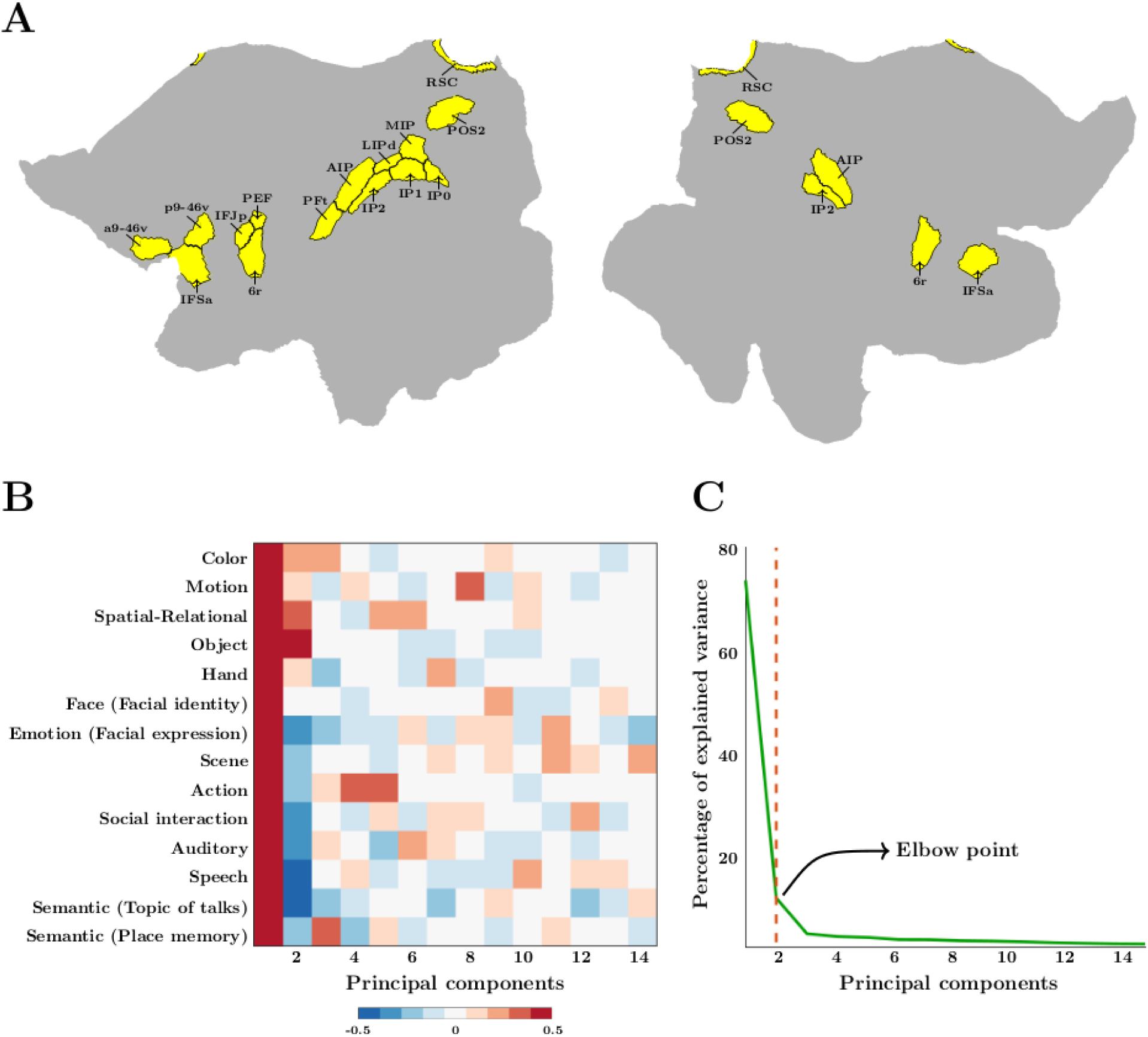
Decomposition of task-specific responses within the MD network. **(A)** Parcel-wise localization of core regions of MD network. Cortical parcels from the HCP MMP with significant task-evoked activation in all 14 cognitive domains relative to the passive-viewing condition are shown. Parcels are displayed on flat surfaces, with the left hemisphere shown on the left and the right hemisphere on the right. **(B)** PCA was applied on task-evoked responses in the MD network. The 14 × 14 matrix shows the correlation coefficients between each of the 14 cognitive tasks (rows) and each of the 14 principal components (columns) derived from PCA. Correlations were computed using activation values (t-values) from all vertices within the core regions of MD network shown in panel A. **(C)** The plot shows the percentage of variance explained by each principal component. The vertical red dashed line marks the elbow point used to determine the number of components retained for subsequent analyses.

### Functional differentiation within the MD network

We next examined whether the MD network exhibits internal specialization across cognitive domains. To this end, principal component analysis (PCA) was applied to vertex-wise task-evoked responses extracted from the core MD network regions/parcels. Task-evoked responses were quantified as t-values derived from group-level contrasts comparing each cognitive task versus passive-viewing condition. This approach identified dominant axes of functional variation within the core MD network and enabled grouping of cognitive tasks based on shared patterns of cortical engagement.

To examine the relationship between the task-evoked activation patterns and their neural representations in the principal component space, we correlated each task-evoked contrast map with principal component scores, yielding a 14 × 14 task–component correlation matrix (Figure 3B). Elbow-point estimation using the Kneedle algorithm [57] indicated that only the first two components accounted for substantial variance (84%) in the data (Figure 3C), justifying their retention for interpretation of task-evoked responses within the MD network.

The first principal component (PC1) exhibited positive correlations across all cognitive domains, indicating that it captures variance shared across all task-evoked activation maps. The magnitude and consistency of these correlations are consistent with a domain-general mode of MD network engagement. The second principal component (PC2) revealed a functional dissociation within the MD network. Tasks that relied on processing/attending/memorizing visual features of the movie clips (i.e., color, motion, spatial-relational, object, hand, and face identity) showed positive correlations. Note that in five of these tasks, the task question was in the pictorial format (Supplementary Figure S1). On the other hand, tasks that required transforming the stimuli into non-visual, categorical, abstract, or semantic representations (i.e., face expression, scene, action, social interaction, auditory, speech, topic of talks, and place memory) showed negative correlations. Note that in all of these tasks, the task question was in the text-based format (Supplementary Figure S1). This pattern suggests that PC2 distinguishes between visually grounded and abstract/semantic modes of cognitive processing within the MD network.

### Spatial organization of functional differentiation within the MD network

Having identified the coexistence of a domain-general and a task-specific component within the MD network, we examined whether this functional differentiation is spatially organized. To address this question, we first visualized the spatial distribution of PC1 scores across MD cortical regions (Figure 4A). The domain-general engagement was not spatially uniform; instead, systematic graded patterns of activation emerged within specific MD subregions. On the lateral parietal surface, the vertices encompassing the positive PC1 scores were concentrated in central portions of LPC, at the borders between posterior-ventral parcels (IP0, IP1, IP2) and anterior-dorsal parcels (MIP, LIPd, AIP) bilaterally. On the medial parietal surface, peaks of positive PC1 scores were observed dorsally within the POS and anteriorly within the RSC. In the frontal cortex, high PC1 values within bilateral inferior frontal junction (IFJ) and inferior frontal sulcus (IFS) were concentrated at the borders between dorsal and ventral parcels, with a greater spatial extent in the left hemisphere relative to the right. Together, these observations indicate that domain-general engagement within the MD network is organized along spatial gradients. These gradients are observed across all patches of the MD network which contribute to all cognitive domains.

**Figure 4:**
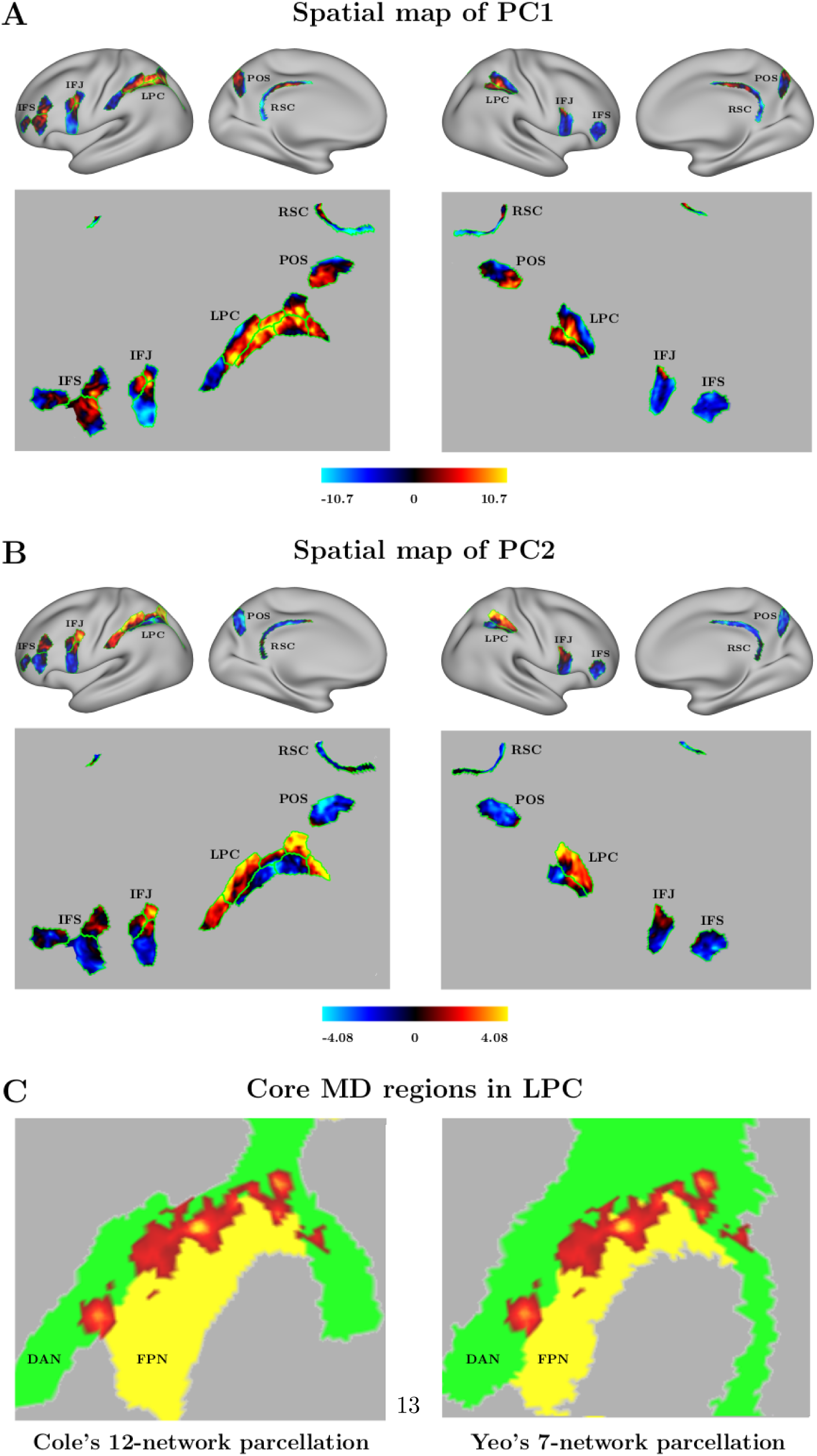
Spatial gradients of principal components within the MD network. Each map shows values of a principal component (PC) across vertices of the MD network. **(A)** Spatial distribution of the first principal component (PC1). **(B)** Spatial distribution of the second principal component (PC2). Borders of the MD network parcels from the HCP MMP are overlaid in green. In panels A and B, left and right hemisphere maps are shown on the left and right, respectively. The top row shows lateral and medial views of an inflated cortical surface, while the bottom row shows flat maps. **(C)** Spatial location of PC1 hotspots (core MD islands) in the left lateral parietal cortex (LPC) relative to the dorsal attention (DAN) and frontoparietal (FPN) networks from Cole’s 12-network parcellation (left) and Yeo’s 7-network parcellation (right).

Beyond this dominant domain-general component, PCA revealed a second component (PC2) that differentiated visual from non-visual/semantic tasks. We next examined whether this functional dissociation exhibited a corresponding spatial pattern by visualizing the spatial distribution of PC2 scores across MD cortical regions (Figure 4B). Within the LPC, which showed the most pronounced gradual changes, positive PC2 values associated with visual task engagement were concentrated in superior regions, whereas negative PC2 values associated with non-visual/semantic task engagement were located more inferiorly. A comparable superior–inferior gradient was evident in the frontal cortex, including IFJ and IFS, with superior portions showing positive PC2 values and inferior portions showing negative PC2 values. In contrast, neither medial parietal region (POS or RSC) exhibited positive PC2 values, indicating a relative bias toward non-visual/semantic processing.

To investigate the organization of domain-general engagement within large-scale cortical networks, we further analyzed the spatial distribution of PC1 values. Prominent peaks of PC1 in the LPC were identified and are hereafter referred to as core MD islands. Notably, these core MD islands were consistently located at the interface between the frontoparietal (FPN) and dorsal attention (DAN) networks, rather than within the interior of either network. This boundary-based organization was preserved across multiple network parcellation schemes [5, 20] (Figure 4C).

We next examined how this pattern of functional specialization relates to surrounding cortical regions by extending the PCA-based approach to the extended MD network (Supplementary Figure 2). The extended MD network was defined as parcels with a probability threshold of *>* 0.5 (i.e., significant activations in more than half of the tasks), then the same PCA analysis was applied to the vertices of these parcels. The resulting maps revealed a broadly similar spatial organization of both PC1 and PC2 across the core and extended MD networks. Notably, extending the analysis beyond the core MD regions clarified the spatial organization of the PC2 component within the frontal cortex. Specifically, vertices selective for visual tasks progressively decreased from superior to inferior frontal regions, accompanied by increasing selectivity for non-visual/semantic tasks in inferior frontal vertices.

Taken together, the spatial distributions of PC1 and PC2 revealed a structured organization within the MD network. PC1, reflecting domain-general engagement, was concentrated centrally within MD regions, forming a core that was gradually surrounded by task-selective components. Relative to this core, PC2 followed a systematic spatial gradient that differentiated visual and non-visual/semantic processing. In lateral regions—including LPC, IFJ, and IFS, visual engagement was preferentially expressed in superior portions, whereas non-visual/semantic engagement was more prominent in inferior portions. This complementary spatial arrangement indicates that domain-general and task-specific processes are not uniformly intermixed but are organized along consistent spatial gradients within the MD network.

### Connectivity signatures of visual–semantic differentiation in the MD network

We next asked whether the observed spatial dissociation between visual and non-visual/semantic components of the MD network (core MD regions) is supported by corresponding differences in intrinsic functional connectivity. Functional connectivity was estimated using an independent movie-watching dataset from the HCP 7T database (https://www.humanconnectome.org/study/hcp-young-adult), as naturalistic viewing provides reliable and information-rich measures of large-scale network organization [58–61]. In this dataset, 176 healthy young adults viewed a series of short audio-visual movie clips (1–4 minutes each) across four runs totaling approximately 60 minutes. The resulting fMRI data were preprocessed, demeaned, concatenated across runs, and averaged across subjects following the procedure described in [61], yielding group-level time-series for connectivity analysis.

Seed regions were defined based on the sign of PC2 values derived from our task-based fMRI dataset. Vertices with positive PC2 scores were assigned to a visual seed, whereas vertices with negative PC2 scores were assigned to a non-visual/semantic seed. For functional connectivity estimation, a single representative time-series was extracted for each seed as a weighted average of the time-series of its constituent vertices, with weights proportional to the absolute PC2 values. Whole-cortex functional connectivity maps were then computed by correlating each seed time-series with the time-series of every cortical vertex.

These seed-based connectivity maps revealed strikingly different patterns for the visual and non-visual/semantic seeds. Visual seed showed strong connectivity with occipital and temporal regions along the ventral visual pathway, as well as with lateral parietal regions within the dorsal visual pathway [62–64] (Figure 5A). Connectivity also extended into the superior parietal cortex, anterior to the MD network, overlapping with regions of the dorsal attention system involved in spatial and feature-based attention [65, 66]. In contrast, semantic seed exhibited a markedly different connectivity profile, showing stronger coupling with lateral prefrontal, inferior parietal, and medial association regions that were largely absent for the visual seed (Figure 5B). To directly compare these patterns, we computed a difference map, contrasting the connectivity of the visual and semantic seed regions (Figure 5C). This map confirmed the dissociation observed in the individual seed-based profiles: visual seed preferentially coupled with perceptual and attention-related systems, whereas semantic seed showed stronger coupling with higher-order association regions. Together, these results indicate that the two PC2-derived seed sets occupy distinct positions within large-scale cortical networks.

**Figure 5:**
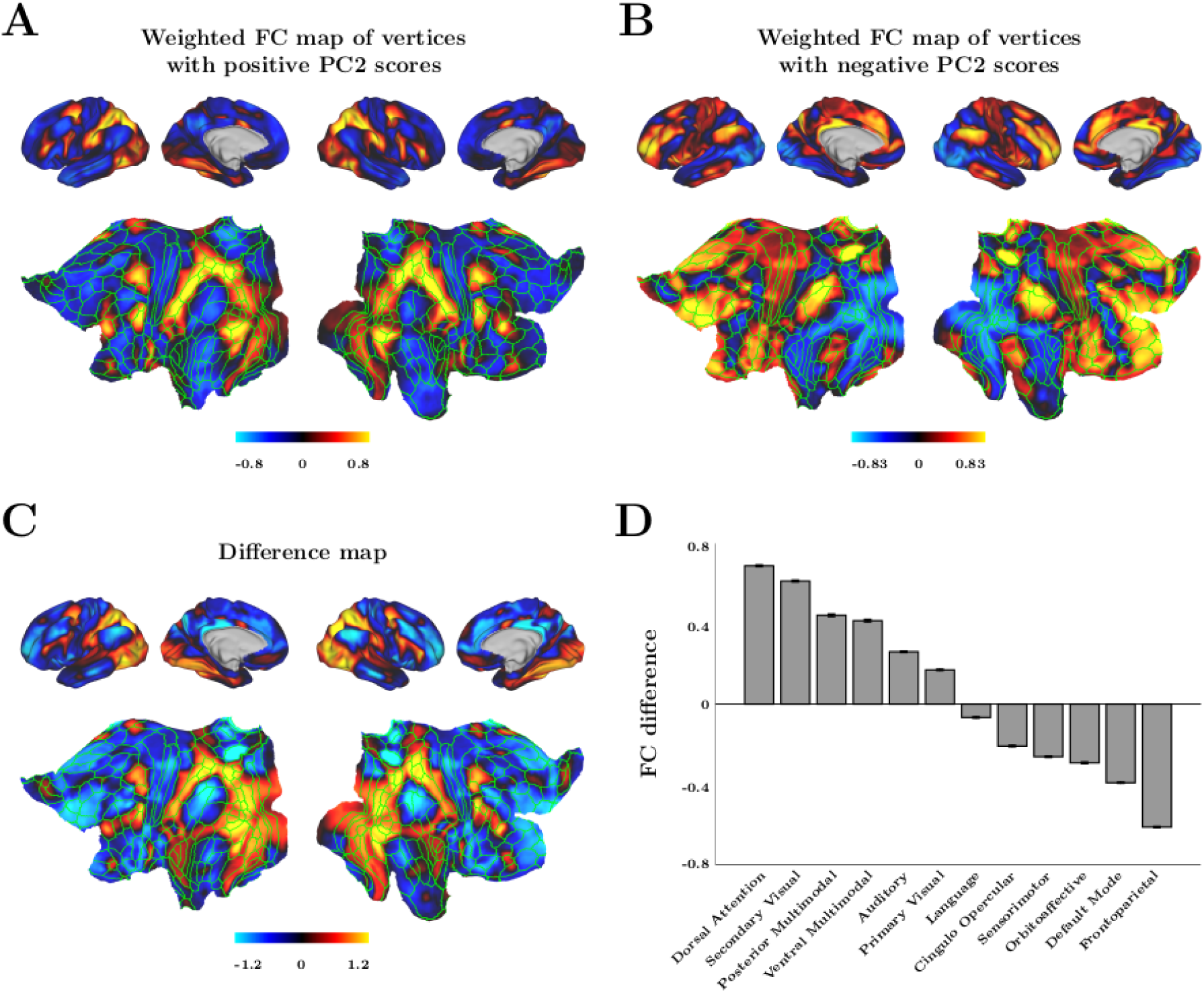
Whole-cortex functional connectivity of visual and non-visual/semantic subdivisions within the MD network. Seed regions were defined based on the sign of PC2 values, yielding a visual seed (vertices with positive PC2 values) and a non-visual/semantic seed (vertices with negative PC2 values). Seed time-series were computed as weighted averages of vertex time-series, with weights proportional to the absolute PC2 values. Time-series were derived from the HCP 7T movie-watching fMRI data averaged across 176 participants. Borders of cortical parcels from the HCP multimodal parcellation (MMP) are overlaid in green. For all maps, left and right hemisphere views are shown on the left and right, respectively. The top row shows lateral and medial views of an inflated cortical surface, and the bottom row shows flat maps. (**A**) Whole-cortex connectivity map for the visual seed. (**B**) Whole-cortex connectivity map for the non-visual/semantic seed. (**C**) Difference map contrasting connectivity of the visual and non-visual/semantic seeds. (**D**) Connectivity differences between the visual and non-visual/semantic seeds across the 12 Cole-Anticevic resting-state networks. Error bars indicate standard error of the mean across vertices within each network.

To further specify these distinctions, we quantified connectivity differences with respect to the 12 canonical resting-state networks defined by Cole and Anticevic [5] (hereafter referred to as the Cole–Anticevic networks) (Figure 5D). The strongest positive differences were observed in the dorsal attention network and secondary visual cortex, reflecting preferential engagement of visual seed with perceptual and attentional systems. In contrast, negative differences were most pronounced in the frontoparietal control and default mode networks, indicating stronger coupling of semantic seed with higher-order cognitive systems.

Together, these findings demonstrate that the functional dissociation captured by PC2 is mirrored in large-scale connectivity architecture: visual regions of the MD network are embedded within perceptual–attentional circuits, whereas semantic regions of the MD network are preferentially integrated with control-related and internally oriented networks.

### Whole-brain graph-theoretic organization of task-related responses

Having established that domain-general and task-differentiated components of the MD network are organized along consistent spatial gradients, we next asked how these components are embedded within the broader architecture of the cortex. To address this question, we examined whole-brain functional organization using graph-theoretic analysis. Rather than limiting the analysis to MD regions, we assessed task-related network structure across the entire cortex. This analysis used cortical parcels defined by the HCP multimodal parcellation, which provides a standardized division of the cortex into 360 parcels (180 per hemisphere).

For each of the 14 task domains, we performed a group-level parcel-wise analysis contrasting task and passive conditions. Each cortical parcel was then characterized by a 14-dimensional response profile, with one value per task domain. Pairwise similarity between parcels was computed using Pearson correlation across these profiles, resulting in a 360 × 360 parcel similarity matrix. To focus on the strongest functional relationships, the matrix was thresholded to retain only high correlations (*r >* 0.8). The resulting adjacency matrix was used to construct a graph in which parcels served as nodes and retained correlations defined edges. The graph was visualized using a force-directed spring-embedding algorithm [67], which positions nodes based on simulated attractive and repulsive forces. For visualization purposes, nodes were colored according to their Cole-Anticevic network identity (Figure 6A). This visualization revealed two clearly separated clusters of nodes, both connected to the frontoparietal/MD network. One cluster comprised parcels from the visual network, including primary and secondary visual cortex, together with nodes from the dorsal attention network. The second cluster was dominated by non-visual networks, including default mode, language, auditory, and sensorimotor systems. Frontoparietal/MD network nodes were distributed between these two clusters. Some MD nodes occupied intermediate positions between the clusters, consistent with a domain-general role, whereas others showed a relative bias toward either the visual or the non-visual cluster.

**Figure 6:**
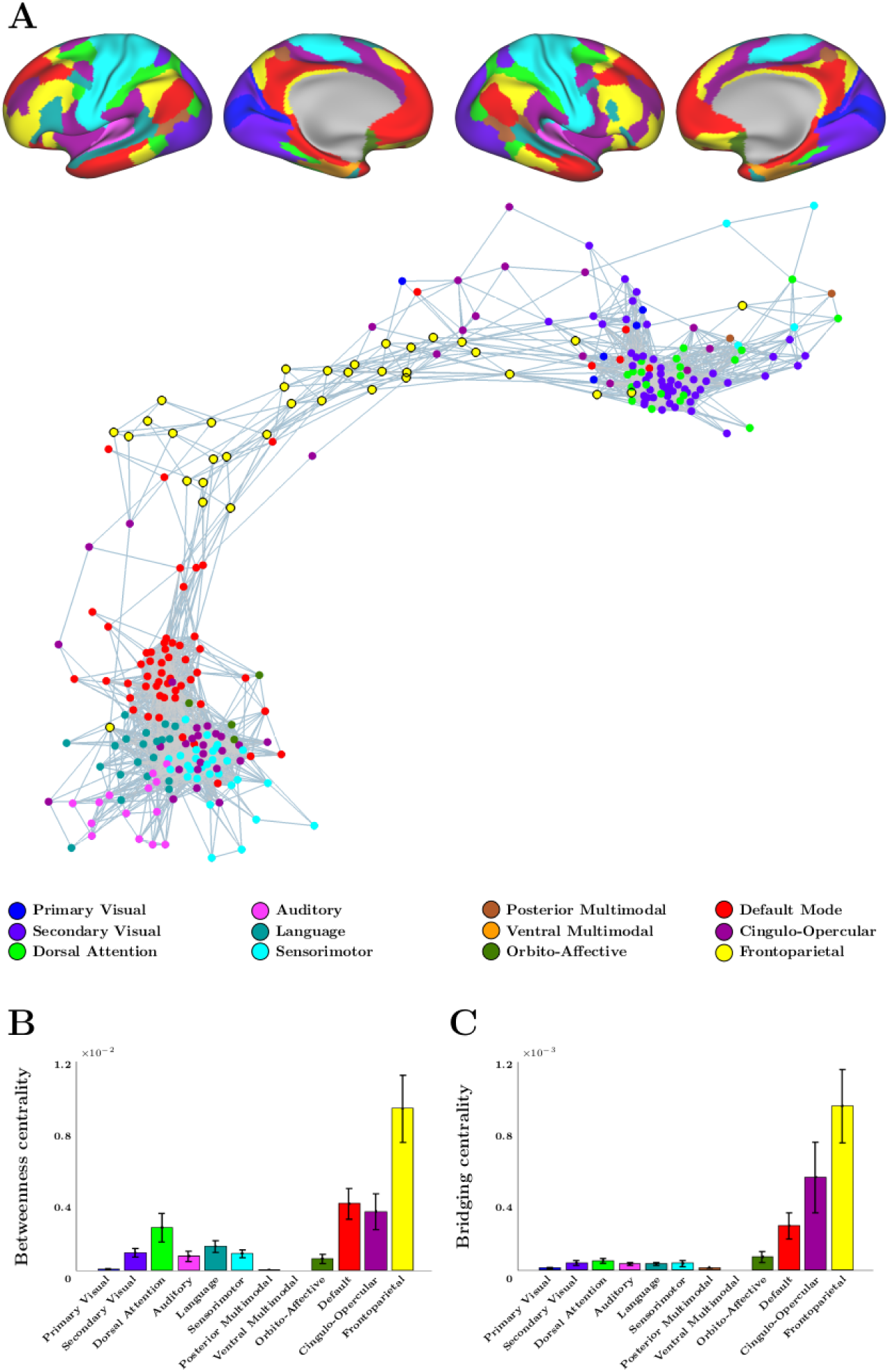
Graph analysis of task-specific activations. (**A**) The top panels show the Cole–Anticevic 12-network parcellation displayed on inflated cortical surfaces of the left and right hemispheres. The bottom panel shows a graph representation of cortical parcel connectivity, derived from parcel-wise task-specific activations across all subjects and the 14 cognitive domains. Graph nodes correspond to parcels from the HCP MMP, with node colors indicating their assigned Cole–Anticevic network. Edges represent pairwise correlations between parcels, thresholded at *r >* 0.8. For visualization, a force-directed spring-embedding algorithm [67] was used, which models edges as springs and iteratively minimizes system energy, such that strongly related parcels cluster while weakly related parcels separate. **(B)** Bar plot of betweenness centrality across cortical parcels, grouped by Cole–Anticevic networks. **(C)** Bar plot of bridging centrality across cortical parcels, grouped by Cole–Anticevic networks. Error bars in panels B and C indicate standard error of the mean across parcels within each network.

To characterize the network topology, we computed two centrality measures (Figure 6B,C). Betweenness centrality quantifies how often a parcel lies on the shortest communication path between other parcels, indexing its contribution to global information transfer [68, 69]. Bridging centrality extends this measure by weighting nodes according to their capacity to connect otherwise segregated communities [70–72]. Both metrics were highest in the frontoparietal and cingulo-opercular networks, which overlap substantially with the core regions of the MD system, confirming their role as dominant hubs linking two large-scale clusters: one comprising visual and attention-related regions and the other encompassing non-visual, higher-order systems. The default mode and dorsal attention networks also showed relatively high betweenness centrality but minimal bridging centrality values, indicating greater involvement in shortest-path communication than in cross-cluster integration. For the dorsal attention network, this pattern aligns with its clustering with visual regions and with our earlier observation that it overlaps primarily with MD network vertices engaged in visual rather than non-visual or semantic processing. In contrast, the default mode network, clustered with non-visual systems, exhibited more localized interconnectivity and a weaker contribution to global integration.

### Behavioral signatures of visual and non-visual task demands

Having identified functional subdivisions within the MD network that differentiate visual and non-visual task demands, we next examined whether a comparable distinction is evident at the behavioral level. Rather than assessing behavior in the same individuals, we analyzed an independent large-scale behavioral dataset to test whether task performance varies systematically across visual and non-visual task demands. The Lumosity platform [73] provides a suitable resource for this purpose. We analyzed behavioral data from 36,297 subjects across 51 cognitive games designed to assess memory, attention, flexibility, speed, problem solving, and decision making, thereby sampling a broad range of cognitive functions. Although all games are visually presented, they vary substantially in the nature of the cognitive operations they require: some emphasize direct visual analysis, whereas others require transforming visual input into abstract, rule-based, or semantic representations.

At the end of each gameplay, users received feedback scores combining accuracy, speed, and bonus points. We quantified behavioral performance as the average total score across gameplays 50–60, chosen to capture stable performance after initial learning curves had plateaued. Performance values were then normalized by scaling each subject’s game scores by their own standard deviation across the final sessions (gameplays 50–60), without mean-centering. This procedure reduced inter-individual differences in response variability while preserving relative performance profiles across tasks. Because subjects did not engage with every game, the resulting task-by-subject matrix was inherently sparse, a limitation further compounded by incomplete learning trajectories in users who discontinued play before reaching 60 gameplays. To extract the dominant dimensions of behavioral variability under these conditions, we applied Bayesian probabilistic PCA [74], which is well suited for sparse datasets and robust to missing observations [75].

The first three dominant components derived from this analysis are shown in Figure 7A. The first principal component (PC1) exhibited uniformly positive loadings across nearly all games, indicating a shared source of variance consistent with general task performance [76]. Consistent with this interpretation, subjects’ PC1 scores were strongly correlated with their mean accuracy across games (*r* = 0.94) (Figure 7B), confirming that PC1 primarily reflects overall behavioral performance.

**Figure 7:**
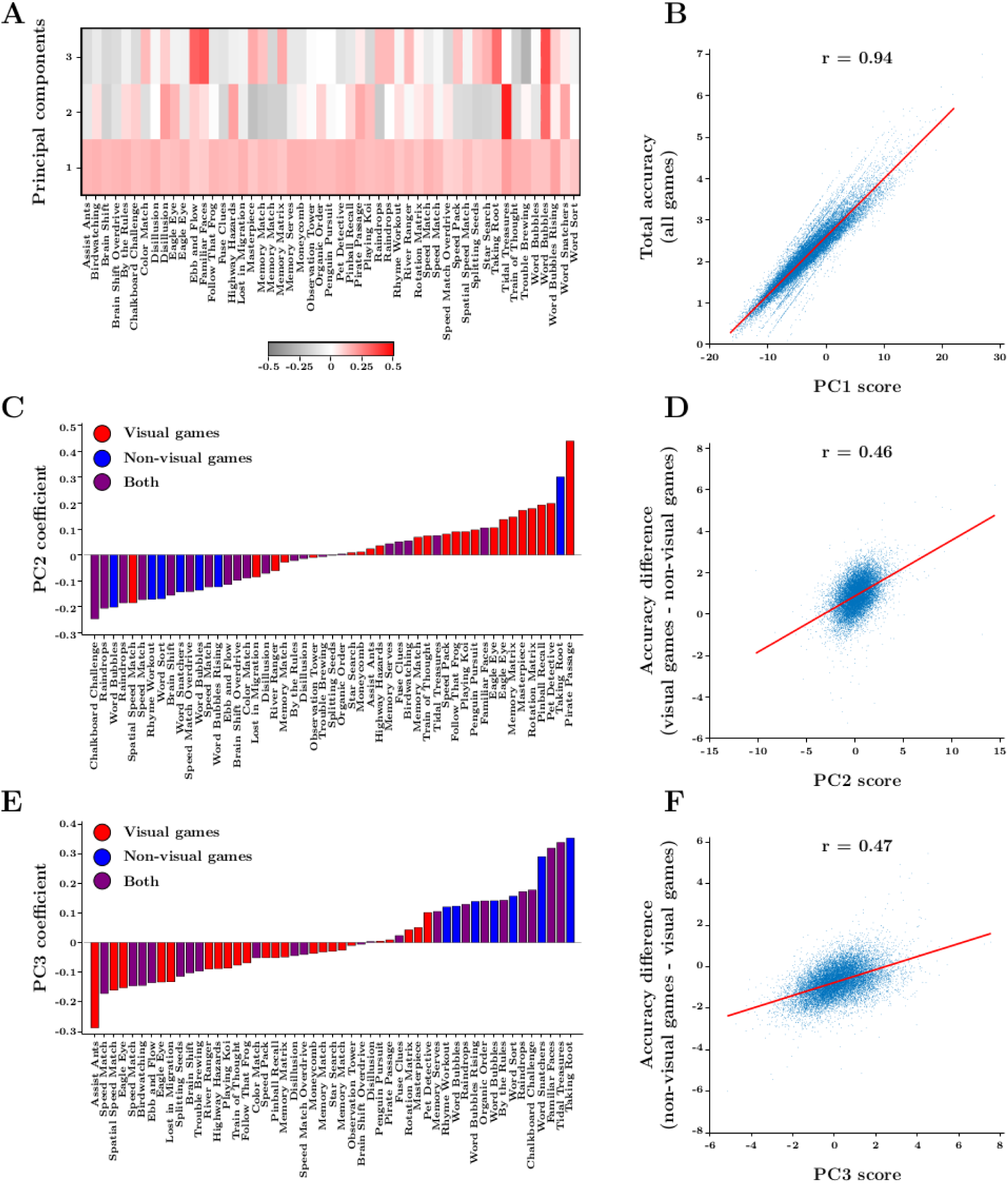
Decomposition of dominant axes of behavioral variation across multiple demanding games. PCA was applied to the performance of subjects across 51 Lumosity games. **(A)** The 51 × 3 matrix of loadings for the first three principal components (PC1–PC3) across all 51 games, derived using probabilistic PCA. Coefficients are color-coded from negative (gray) to positive (red) and clipped to the range of –0.5 to 0.5. **(B)** Scatter plot of subjects’ total accuracy across games (y-axis) versus PC1 scores (x-axis). A fitted regression line is shown in red, revealing a strong positive correlation (r = 0.94), indicating that PC1 primarily reflects overall task performance. **(C)** Bar plot of PC2 loadings for each game, sorted by PC2 coefficient. Visual, non-visual, and mixed games are shown in red, blue, and purple, respectively. Games transition from predominantly non-visual at low PC2 values to predominantly visual at high PC2 values, revealing a clear visual vs. non-visual task gradient. **(D)** Scatter plot of subjects’ PC2 scores (x-axis) versus their accuracy difference between visual and non-visual games (y-axis). A fitted regression line is shown in red (r = 0.46), indicating that PC2 captures individual differences in domain-specific performance across visual and non-visual domains. **(E)** Bar plot of PC3 loadings for each game, sorted by PC3 coefficient. Visual, non-visual, and mixed games are shown in red, blue, and purple, respectively. Games transition from predominantly visual at low PC3 values to predominantly non-visual at high PC3 values, revealing a second task-type gradient. **(F)** Scatter plot of subjects’ PC3 scores (x-axis) versus their accuracy difference between non-visual and visual games (y-axis). A fitted regression line is shown in red (r = 0.47), indicating that PC3 captures another dimension of individual differences in domain-specific performance.

To assess whether behavioral variability beyond the general performance component relates to differences in task demands, we next examined whether the visual vs. non-visual distinction was expressed along subsequent principal components. Games were categorized as visual or non-visual based on descriptions provided in the Lumosity Cognitive Training (CT) documentation, which details the cognitive demands and task characteristics of each game [77]. These descriptions were independently evaluated by four large language models: ChatGPT (GPT-4o family, OpenAI), Gemini (Gemini 1.5 Flash/Pro, Google), Claude (Claude 3.5 Sonnet, Anthropic), and DeepSeek (DeepSeek V2/V2.5 series, DeepSeek AI). Each model assigned games to visual or non-visual categories based solely on the provided task descriptions. Classifications were then aggregated to assign each game to one of three final categories (visual, non-visual, or both), thereby minimizing subjective interpretation and reducing potential model-specific bias. In cases where the models disagreed, the game was considered both visual and non-visual.

The second principal component (PC2) revealed a clear separation between visual and non-visual tasks (Figure 7C). When tasks were sorted by their PC2 loadings, games transitioned smoothly from predominantly non-visual tasks with negative coefficients to predominantly visual tasks with positive coefficients, forming a continuous task-type gradient. This pattern indicates that PC2 captures systematic differences in the cognitive demands of visual versus non-visual processing. Consistent with this interpretation, subjects’ PC2 scores were positively correlated with their accuracy difference between visual and non-visual games (*r* = 0.46) (Figure 7D), demonstrating that PC2 reflects individual differences in task-specific behavioral performance.

The third principal component (PC3) displayed a similar gradient but in the opposite direction (Figure 7E). Games primarily engaging visual processing showed negative loadings, whereas those involving non-visual demands had positive loadings. This axis therefore captures a complementary dimension of the behavioral pattern, differentiating visual and non-visual task engagement along a secondary source of variance. Subjects’ PC3 scores were correlated with their accuracy difference between non-visual and visual tasks (*r* = 0.47) (Figure 7F), further confirming that PC3 represents an additional behavioral component related to individual differences in task-specific performance.

## Discussion

A central question in the MD-network literature is whether this system operates as a largely homogeneous, domain-general controller or whether it contains meaningful internal specialization. We addressed this question using a naturalistic paradigm in which sensory input was held constant while participants engaged in a broad set of cognitive tasks, allowing us to characterize the organization of the MD network across diverse task demands. Our results show that the MD network comprises a domain-general core alongside task-differentiated components expressed across functional activation, spatial organization, functional connectivity, and behavior, with visual versus non-visual control demands emerging as a primary axis shaping this subdivision.

In the present study, the MD network was localized using naturalistic movie stimuli that engaged a broad range of cognitive domains, yielding a spatial pattern consistent with previous reports. Earlier work has identified the MD network using more restricted sets of modality-specific tasks [17] or through aggregation across meta-analytic contrasts [78, 79]. Our approach, however, provided a more integrated characterization by systematically varying cognitive demands while holding sensory input constant. This design combines ecological validity with experimental control, enabling domain-general and task-differentiated contributions within the MD network to be assessed within a single unified framework. The close correspondence between our MD maps and HCP-based MD maps [23] further supports the validity of this approach and highlights the utility of naturalistic paradigms for identifying domain-general control networks.

Within the localized MD network, our differentiation analysis supports a hybrid model of MD organization, balancing domain-general and task-specific components. First, we observed a dominant component expressed across all tasks, concentrated in canonical frontal and parietal MD territories, aligning with longstanding hypotheses that the MD network functions as a domain-general control system recruited by diverse cognitive demands [8, 9, 23]. Second, we identified systematic task-differentiated structure: a visual versus non-visual/semantic axis accounted for additional variance in the MD responses. This finding challenges both strictly homogeneous models and those that partition the MD network into discrete executive function modules [17, 47, 50]. Instead, task differentiation was expressed as smooth spatial gradients within established MD territories, with transitions from task-general peaks to more task-differentiated responses occurring without sharp boundaries. This organization is consistent with accounts proposing that domain-general cores are embedded within broader cortical territories whose functional profiles are shaped by task demands and connectivity to neighboring systems [53–55], thereby reconciling domain-general and task-differentiated views of MD function.

One prominent feature of our spatial localization results is that the domain-general component, particularly within the LPC patch, peaked at the boundary between FPN and DAN, rather than within the interior of a single network. This pattern closely mirrors recent evidence showing that the strongest expressions of executive control responses are displaced toward transitions between the MD network and other resting-state networks, with task switching exhibiting a pronounced bias toward DAN-adjacent boundaries [50]. These observations build on a broader literature proposing that regions positioned between large-scale networks, often described as ’connector hubs’, are especially well suited for integrating information across systems [80, 81]. Rather than belonging exclusively to a single functional network, such regions appear to derive their importance from their intermediate embedding within the cortical architecture. From this perspective, the border-peaked MD activations observed here may reflect locations where domain-general control processes are optimally positioned to interface with other systems, supporting flexible coordination across diverse task demands.

A key finding of the present study is a dorsal–ventral organization within the MD network that tracks a visual to non-visual/semantic axis of control demands. Rather than forming sharply bounded subregions, MD responses varied continuously along this axis: tasks relying more directly on visually grounded processing preferentially engaged dorsal MD territories, whereas tasks requiring transformation into non-visual, semantic representations preferentially engaged ventral MD territories. This gradient-like organization aligns with established principles of cortical architecture in which large-scale functional gradients extend from sensory cortex toward increasingly sensory-motor distal association regions [82, 83]. Within this framework, frontoparietal/MD control regions occupy intermediate positions, supporting smooth transitions in the form of control engaged as task demands shift from perceptually anchored operations toward more semantically and conceptually guided transformations, consistent with prior accounts of dorsal–ventral differentiation in the frontoparietal control system [84–88].

Viewed from a larger-scale systems perspective, functional connectivity provides a plausible architectural substrate for this graded organization. MD territories aligned with the visually oriented end of the organizational axis were more strongly coupled with occipital and ventral temporal visual cortex, as well as dorsal parietal regions associated with visual processing and spatial attention, whereas MD territories aligned with the non-visual end of the axis showed stronger coupling with lateral prefrontal and medial association regions implicated in semantic and internally oriented processing [62, 65, 66, 89]. Converging with this dissociation, whole-brain graph analyses revealed two corresponding large-scale configurations: one in which frontoparietal/MD regions interface with visual networks via dorsal attention system, and another in which MD regions interface with non-visual association networks via default-mode circuitry [5, 21, 53]. This pattern is consistent with the idea that distinct MD subterritories maintain relatively stable affiliations with different large-scale networks, providing a systems-level basis for flexible recruitment across heterogeneous task demands [52, 53].

Notably, the visual–conceptual differentiation observed within the MD system echoes a broader organizational motif seen across category-selective pathways. In scene-, face-, and object-selective systems, posterior territories tend to be driven more strongly by perceptual features, whereas progressively anterior regions support increasingly conceptual, mnemonic, or semantic representations [90–93]. The presence of an analogous distinction within a domain-general control network suggests that the MD system may rely on a heterogeneous representational architecture to control distinct forms of information. In this view, the MD system may flexibly coordinate perceptual input and higher-order transformations within a unified control framework by drawing on subterritories differentially embedded within sensory-attentional versus association-level systems, rather than relying on a single undifferentiated control mechanism [5, 23].

Behavioral data provide a complementary perspective on the visual and non-visual/semantic differentiation observed in the MD network. In an independent large-scale cognitive dataset, inter-individual variability in task performance decomposed into a dominant domain-general factor alongside secondary and tertiary axes differentiating visually grounded from more abstract, non-visual task demands. Behavioral performance exhibited a hierarchical organization, such that at the most general level, performance was positively correlated across domains, consistent with the positive manifold, the well-established tendency for cognitive abilities to covary across individuals [94, 95]. This domain-general performance factor aligns with prior neuroimaging work demonstrating that activity within the frontoparietal MD network scales with individual differences in general intelligence and executive control [13, 96, 97]. Beyond this shared component, performance decomposed into orthogonal dimensions reflecting modality-specific specialization: some individuals showed relative strengths in visually grounded perceptual tasks, whereas others were more proficient in abstract or semantic domains. Converging recent evidence suggests that such performance profiles reflect stable differences in cognitive strategy and representational format, with individuals differing in the extent to which they rely on visual versus non-visual, abstract control processes during complex cognition [13, 32, 98]. Importantly, these strategy-linked modality preferences did not reflect a trade-off in overall ability, as performance in visual and non-visual tasks remained positively associated. Together, these findings suggest that individual differences in behavior may arise not only from variation in overall cognitive capacity, but also from systematic differences in how functionally distinct subcomponents of the MD network are differentially engaged or weighted across individuals.

However, while the pattern of these behavioral differences resembles the functional organization we observed in the MD network, it is important to recognize that the two datasets are not directly coupled. In the neuroimaging component of our study, we do not have access to detailed behavioral profiles that could clarify individual-level specialization. Conversely, in the behavioral dataset, neuroimaging data are not available to support direct mapping onto specific neural substrates. As such, while the correspondence between neural and behavioral patterns is suggestive, it should not be interpreted as a one-to-one relationship. Furthermore, alternative explanations may account for the observed behavioral distinctions. For instance, prior studies have distinguished between ’fluid intelligence’ linked to flexible problem solving and reasoning and ’crystallized intelligence’ which reflects knowledge accumulated through learning and experience [99, 100]. The MD network has been repeatedly implicated in fluid intelligence [8, 31], whereas crystallized intelligence may rely more on temporoparietal and semantic processing systems [101]. It is therefore plausible that individual differences in domain preference reflect differing balances between fluid and crystallized cognitive abilities, rather than or in addition to fine-grained organization within the MD system.

Together, these considerations emphasize the need for future studies that integrate behavioral and neural data within the same individuals to more directly assess how large-scale brain networks support different cognitive profiles. Nonetheless, the convergence observed here across independent datasets provides preliminary support for a model in which flexible cognition arises from the interplay of domain-general executive mechanisms and individual differences in cognitive specialization.

## Methods

### Subjects

In this study, a total of 20 subjects (11 female, 9 male, aged 22-34) were scanned. All subjects demonstrated normal vision, normal auditory function, and good English listening comprehension. Subjects were recruited through advertisements, were compensated for their time, and provided written informed consent. The experimental protocol was approved by the ethics committee at the Institute for Research in Fundamental Sciences (approval number: SCS. REC: 1402/40/1/4817).

### Stimuli

The experimental stimuli consisted of specially designed audio–visual movie clips, each constructed to incorporate all fourteen cognitive domains (see Figure 1B for the full list). The movie clips were adapted and edited from Hollywood films to ensure suitability for the experimental paradigm and to provide a balanced representation of perceptual, emotional, social, and semantic content. Each movie clip included dynamic, colorful, and naturalistic scenes combining both indoor and outdoor environments with rich motion, object interactions, social exchanges, and conversations. Each movie clip contained at least two social interactions featuring a mix of familiar and unfamiliar characters, with clear variations in facial expressions to capture emotional content and sustained visual attention to faces. Scenes incorporated recognizable landmarks, flags, or other contextual cues to provide spatial and semantic references. To ensure that information from all domains was evenly represented, segments from different films were occasionally merged, with a continuous soundtrack applied to maintain coherence and prevent abrupt transitions. Soundtracks were equalized in volume to ensure perceptual consistency, and non-vocal natural sounds [102] relevant to the scenes were blended to enhance ecological validity and provide semantically meaningful auditory cues. In total, fourteen movie clips were created: two were reserved for practice, and the remaining twelve were randomly divided into two sets of six movies for use in the passive-viewing and task-based fMRI runs, with the assignment of sets varying across subjects.

### Experimental paradigm

Data were collected in three sessions. The first session included structural, field mapping, resting-state, and passive-viewing scans. Resting-state fMRI data were acquired in two runs (∼ 7.5 min each) while subjects rested with eyes closed. This was followed by two passive-viewing runs (∼ 5 min each), during which subjects viewed twelve experimental movie clips presented across two complementary sets of six. Each run followed a standardized trial structure comprising a cue–movie–question sequence (Figure 1A): a 1-second visual cue (Just watch the movie!) appeared at the center of the screen, followed by a movie clip (26–37 s) and a two-choice arithmetic question (e.g., simple single- or two-digit addition) unrelated to the movie content, to which subjects selected the correct answer. This procedure established the standard timing, perceptual (visual and auditory), and motor requirements applied in all subsequent runs (practice and task-based), while avoiding domain-specific attentional demands. Following the first session, subjects received an instructional pamphlet outlining the structure and attentional focus of the upcoming task-based runs. The pamphlet summarized the cue categories and specified the type of information to attend to in each condition, using concise pictorial and textual examples to illustrate task instructions.

The second and third sessions comprised the main task-based fMRI runs and field mapping scans. Prior to scanning, subjects completed four practice runs (∼ 5 min each) outside the scanner to gain practical familiarity with the question–response format and attentional focus introduced in the instructional pamphlet. These runs covered all fourteen cognitive domains but used different movies and questions from those presented during fMRI acquisition, while maintaining the same trial structure (cue–movie–question sequence). This ensured procedural familiarity while avoiding overlap between practice and experimental stimuli. After completing the practice runs, subjects performed fourteen task-based fMRI runs (∼ 6 min each) in each session. Each run targeted one cognitive domain and consisted of six trials, following the same trial structure described for the passive-viewing runs. Here, however, the cue indicated the target domain (e.g., color, motion, emotion, or speech), which subjects were instructed to attend to selectively during the subsequent movie clip. Cues were presented in the same format across all trials within a run. For the color and spatial–relational conditions, additional subcues were introduced to balance attentional demands and simplify task interpretation. In the color condition, subcues specified the color of dresses, objects, or scenes. In the spatial–relational condition, they specified person–object, person–person, or person–scene relations. Across runs, the same set of six movies was presented in randomized order to ensure consistent stimulus exposure while minimizing sequence-related confounds.

After each movie, subjects answered a two-choice question directly related to the cued domain. For verbal domains (emotion, scene, action, social interaction, auditory, speech, and semantics), questions were presented as true/false statements or sentence completions, and subjects selected the correct or most context-appropriate textual option. For visual domains (color, motion, object, body, and face), questions were in a visual two-choice format in which subjects viewed two images and identified the one corresponding to the movie content. The images used for all visual domains were presented in grayscale and matched in size, except for the color condition, where the images were colorful frames extracted from the corresponding movie frames to preserve color information relevant to the task. Responses were made via button press within a 10-second window, followed by a blank screen for the remainder of the trial.

### Data acquisition

MRI data acquisition followed the core framework of the Human Connectome Project (HCP) 3T protocol [103,104], adapted for the hardware and experimental constraints of the National Brain Mapping Laboratory (NBML) in Iran. Data were collected using a 3T Siemens Magnetom Prisma scanner equipped with a standard 20-channel RF receive head coil.

One 3D T1w MPRAGE image and one 3D T2w SPACE image were collected at 1 mm isotropic resolution.

Functional data of resting-state, passive-viewing, and task-based scans were collected using a simultaneous multi-slice echo planar imaging (SMS EPI) sequence with the following parameters: repetition time (TR) = 1200 ms, echo time (TE) = 30 ms, flip angle = 80 deg, matrix = 64 × 64, number of slices = 42, voxel size = 3 × 3 × 3 mm^3^ isotropic, multiband factor = 2, GRAPPA acceleration factor = 2. The direction of phase encoding alternated between posterior-to-anterior (PA) and anterior-to-posterior (AP) across runs. Gradient-echo field maps were collected during functional MRI sessions to correct spatial distortions in fMRI caused by magnetic field inhomogeneities.

### Data analysis software

The software packages used for analysis included FMRIB Software Library (FSL) [105], FSL ICA-based Xnoiseifier (FIX) [106,107], FreeSurfer [108], Connectome Workbench commandline tools and GUI ’wb view’ [109], MATLAB, and R packages.

### Data analysis

Data were minimally preprocessed and analyzed using the Human Connectome Project (HCP) pipelines [104], version 4.7.0 (https://github.com/Washington-University/HCPpipelines).

#### Structural MRI preprocessing and analysis

Structural MRI data were first preprocessed in volume space using FSL applications, followed by surface reconstruction and subcortical segmentation performed with FreeSurfer.

In volume space, T1- and T2-weighted images underwent brain extraction, correction for intensity inhomogeneity, and cross-modal intensity normalization. The T2w volume was rigidly registered to the corresponding T1w volume, which was subsequently nonlinearly aligned to the MNI152 template. These procedures ensured accurate anatomical correspondence across modalities and subjects. Subcortical gray matter structures were then delineated to provide anatomical constraints for cortical surface reconstruction.

In surface space, the white matter surface was reconstructed along the gray–white boundary constrained by the T1-weighted contrast, and the pial surface was generated along the gray–CSF boundary guided by the T2-weighted contrast. The cortical ribbon, representing the gray matter sheet between the white and pial surfaces, was extracted to define the cortical gray matter. In a separate step, T1w/T2w-based myelin maps were calculated to quantify cortical microstruc-ture [110, 111]. Cortical surfaces were resampled to the fsaverage mesh and then to the fs LR 32k (Conte69) mesh for intersubject comparison. Surface registration was initially based on cortical folding geometry (MSMSulc) and later refined using multimodal surface matching (MSMAll) algorithm, which integrated structural features such as sulcal depth and myelin maps with functional features such as resting-state networks and visuotopy to improve cross-subject correspondence [112].

#### Functional MRI preprocessing

In volume space, preprocessing included motion correction, distortion correction of EPI images using gradient echo field maps, registration of EPI images to the structural T1-weighted scan, nonlinear registration of the native volume to MNI space, brain extraction, and grand-mean intensity normalization. The volume data in MNI space were then segmented to extract subcortical gray matter structures and were smoothed using kernels constrained by the boundaries of these segmented structures.

In surface space, functional data were first projected from volume to the cortical ribbon using a partial-volume–weighted, ribbon-constrained mapping algorithm. Resting-state data were then preprocessed to use in the MSMAll registration and ICA+FIX denoising (component classification). Once the MSMAll registration was established, all functional datasets (resting-state, passive-viewing, and task-based runs) were resampled to the MSMAll-aligned fs LR 32k space and minimally smoothed on the surface using a 2D Gaussian kernel with FWHM = 2. Finally, cortical time-series from both hemispheres and parcel-constrained subcortical voxel time-series were combined into a single grayordinates CIFTI time-series for subsequent analyses.

#### Estimation of task-evoked activations

In each cortical vertex of the preprocessed passive-viewing and task-based CIFTI time-series, task-evoked activity was estimated using a first-level general linear model (GLM). Prior to model estimation, time-series were high-pass filtered to remove low-frequency drift and whitened to account for temporal autocorrelation. For all runs, the GLM design matrix included regressors modeling the cue, movie, question, response, and blank event periods. Regressors were generated by convolving boxcar event functions with a double-gamma hemodynamic response function (HRF) and its temporal derivative. For each run, a contrast of parameter estimates (COPE) was computed for the movie-versus-blank comparison. Within each subject, COPE maps from the two runs of each condition were combined using fixed-effects averaging.

At the group level, we conducted a random-effects analysis contrasting task-based and passive-viewing conditions at both vertex-wise and parcel-wise levels. At the vertex level, paired-sample t-tests were performed at each vertex to compare activation between conditions, yielding vertex-wise t- and p-values. At the parcel level, COPE values from the movie-versus-blank contrast were averaged across vertices within each parcel and then entered into a group-level paired-sample t-test, yielding parcel-wise t- and p-values.

## Data availability

The collected imaging data used in this manuscript are available from the authors upon reasonable request. The open-access HCP data are available at https://www.humanconnectome.org. The Lumosity dataset is available at https://osf.io/g9zkf/.

## Code availability

All analysis codes and scripts are available for sharing upon request.

## Acknowledgments

We thank technical staff of NBML in Iran for help with data acquisition, and Robert J. Schafer for introducing the Lumosity dataset. This research was supported by Iranian Institute for Research in Fundamental Sciences (IPM).

## Author contributions

R.R. conceived the experiment and planned the analyses; N.A. collected the data; M.E.K. contributed to the data collection; N.A. analyzed the data and prepared the figures; N.A. wrote the manuscript; R.R. critically revised the manuscript. All authors approved the final version of the manuscript.

## Competing interests

The authors declare no competing interests.

## Supplementary figures

**Supplementary Figure 1:**
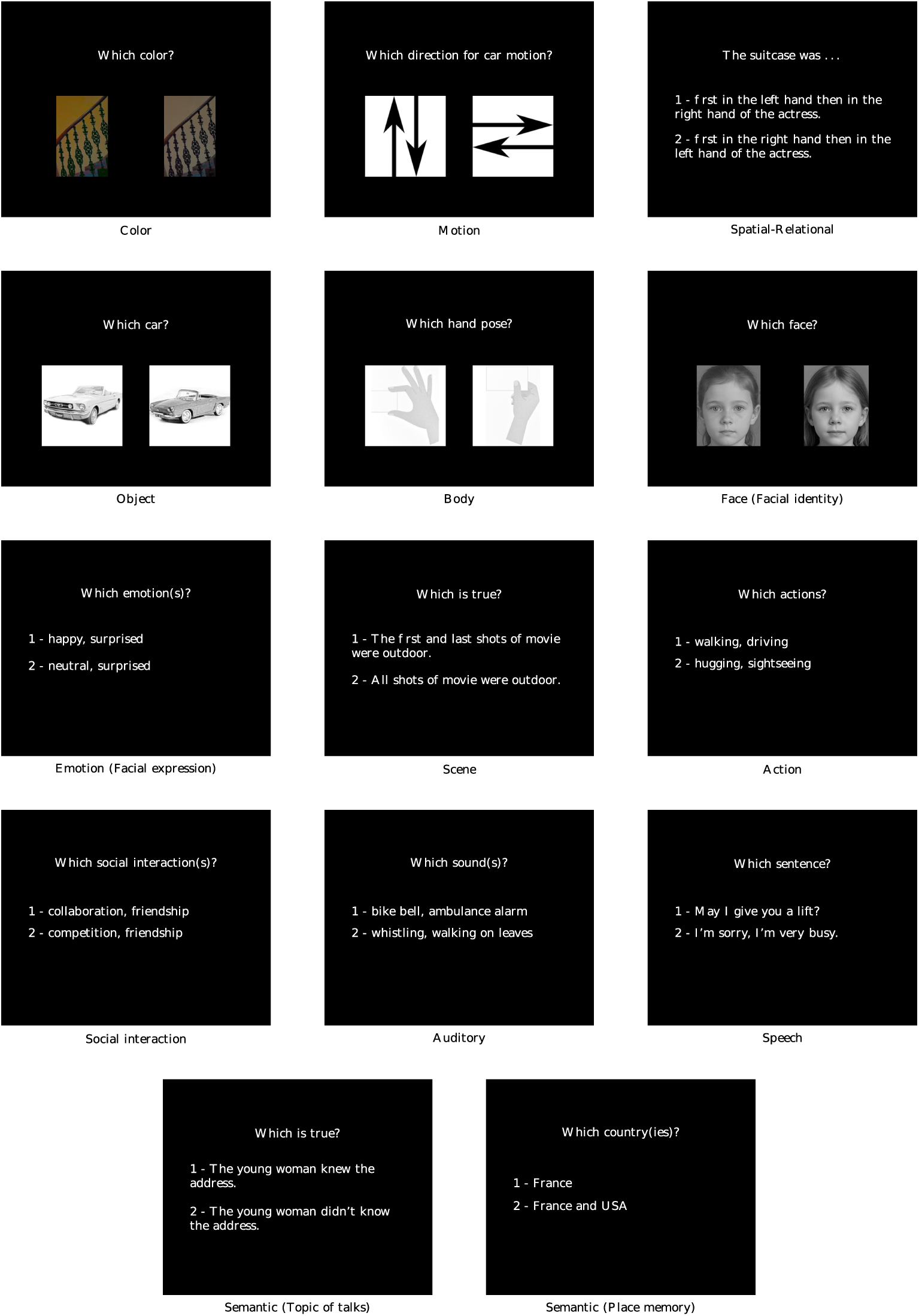
Representative examples of the question and answer presented during the question event in the task-specific runs, for all cognitive domains. The two-choice questions were designed to probe domain-specific attention, and included visual formats (image-based choices) and verbal formats (true/false statements or sentence completions). In the face (facial identity) condition, the original face stimuli used in the experiment are not shown due to copyright restrictions and have been replaced with AI-generated faces for illustrative purposes.

**Supplementary Figure 2:**
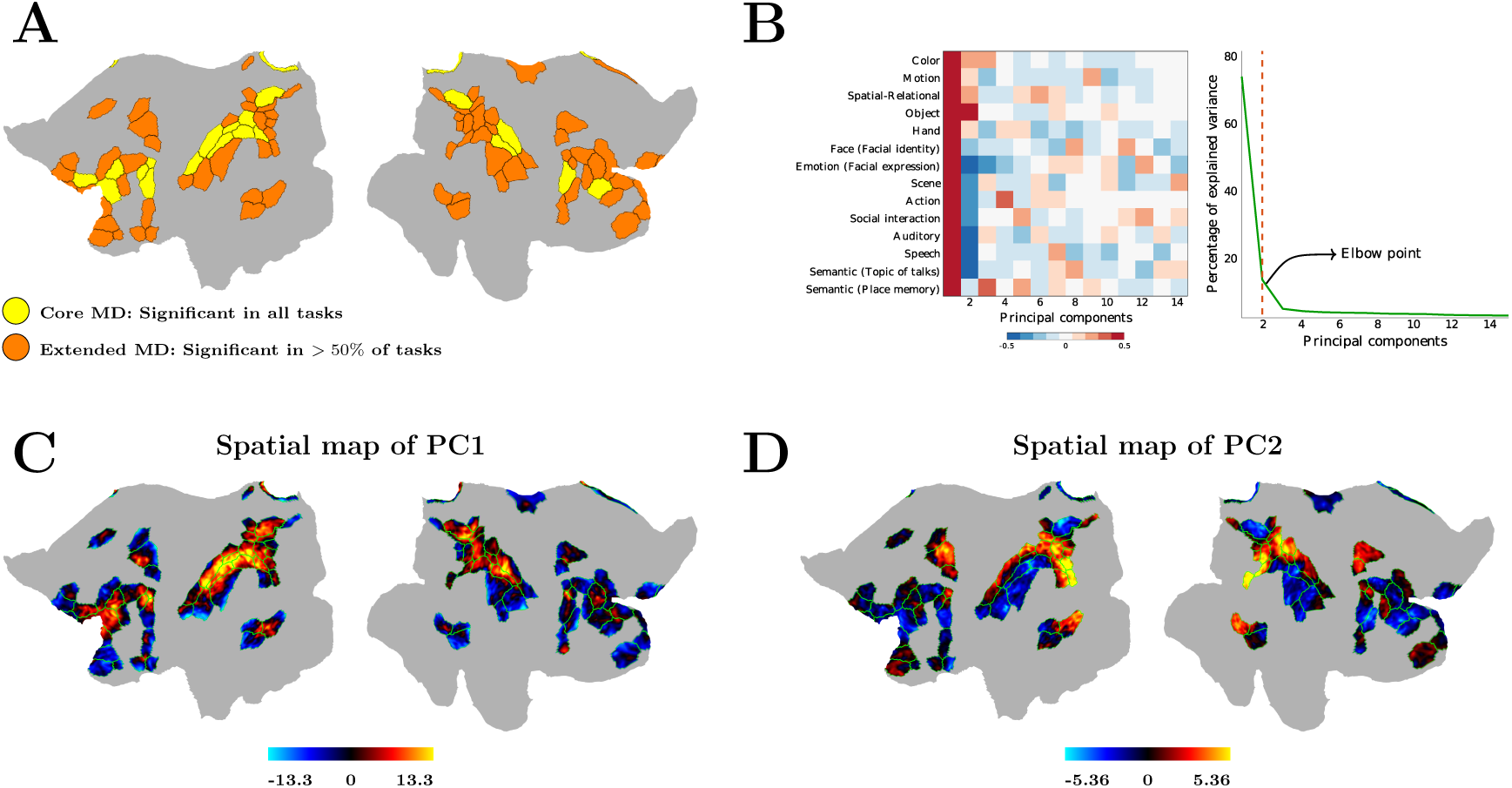
Decomposition of task-specific responses within the extended MD network. **(A)** Cortical parcels from the HCP MMP with significant task-evoked activation in at least 50% of the cognitive domains relative to the passive-viewing condition are shown in orange (extended MD regions), whereas parcels significantly activated in all cognitive domains are shown in yellow (core MD regions). **(B)** PCA was applied on task-evoked responses in the extended MD network (extended and core MD regions). *Left:* The 14 × 14 matrix shows the correlation coefficients between each of the 14 cognitive tasks (rows) and each of the 14 principal components (columns) derived from PCA. Correlations were computed using activation values (t-values) from all vertices within the extended MD network shown in panel A. *Right:* The plot shows the percentage of variance explained by each principal component. The vertical red dashed line marks the elbow point used to determine the number of components retained for subsequent analyses. **(C)** Spatial distribution of the first principal component (PC1) across vertices of the extended MD network. **(D)** Spatial distribution of the second principal component (PC2) across vertices of the extended MD network. Borders of the MD parcels are overlaid in green. In panels A, C, and D, flat maps of left and right hemispheres are shown on the left and right, respectively.

## Notes

### Competing Interest Statement

The authors have declared no competing interest.

## References

[1] Miller, E. K., & Cohen, J. D. (2001). An integrative theory of prefrontal cortex function. Annual Review of Neuroscience, 24, 167–202. 10.1146/annurev.neuro.24.1.167

[2] Egner, T., & Liu, A. S. (2024). Insights into control over cognitive flexibility from studies of task-switching. Current Opinion in Behavioral Sciences, 55, 101342. 10.1016/j.cobeha.2023.101342

[3] Shenhav, A., Musslick, S., Lieder, F., Kool, W., Griffiths, T. L., Cohen, J. D., & Botvinick, M. M. (2017). Toward a rational and mechanistic account of mental effort. Annual Review of Neuroscience, 40, 99–124. 10.1146/annurev-neuro-072116-031526

[4] Cole, M. W., Yarkoni, T., Repov̌s, G., Anticevic, A., & Braver, T. S. (2012). Global connectivity of prefrontal cortex predicts cognitive control and intelligence. The Journal of Neuroscience, 32(26), 8988–8999. 10.1523/JNEUROSCI.0536-12.2012

[5] Cole, M. W., Reynolds, J. R., Power, J. D., Repov̌s, G., Anticevic, A., & Braver, T. S. (2013). Multi-task connectivity reveals flexible hubs for adaptive task control. Nature Neuroscience, 16(9), 1348–1355. 10.1038/nn.3470

[6] Barbey, A. K. (2018). Network neuroscience theory of human intelligence. Trends in Cognitive Sciences, 22(1), 8–20. 10.1016/j.tics.2017.10.001

[7] Duncan, J., & Owen, A. M. (2000). Common regions of the human frontal lobe recruited by diverse cognitive demands. Trends in Neurosciences, 23(10), 475–483. 10.1016/S0166-2236(00)01633-7

[8] Duncan, J. (2010). The multiple-demand (MD) system of the primate brain: Mental programs for intelligent behaviour. Trends in Cognitive Sciences, 14(4), 172–179. 10.1016/j.tics.2010.01.004

[9] Fedorenko, E., Duncan, J., & Kanwisher, N. (2013). Broad domain generality in focal regions of frontal and parietal cortex. Proceedings of the National Academy of Sciences, 110(41), 16616–16621. 10.1073/pnas.1315235110

[10] Fox, M. D., Snyder, A. Z., Vincent, J. L., Corbetta, M., Van Essen, D. C., & Raichle, M. E. (2005). The human brain is intrinsically organized into dynamic, anticorrelated functional networks. Proceedings of the National Academy of Sciences of the United States of America, 102(27), 9673–9678. 10.1073/pnas.0504136102

[11] Duncan, J. (2001). An adaptive coding model of neural function in prefrontal cortex. Nature Reviews Neuroscience, 2(11), 820–829. 10.1038/35097575

[12] Naghavi, H. R., & Nyberg, L. (2005). Common fronto-parietal activity in attention, memory, and consciousness: shared demands on integration?. Consciousness and cognition, 14(2), 390–425. 10.1016/j.concog.2004.10.003

[13] Assem, M., Blank, I. A., Mineroff, Z., Ademoglu, A., & Fedorenko, E. (2020). Activity in the fronto-parietal multiple-demand network is robustly associated with individual differences in working memory and fluid intelligence. Cortex, 131, 1–16. 10.1016/j.cortex.2020.06.013

[14] Rottschy, C., Langner, R., Dogan, I., Reetz, K., Laird, A. R., Schulz, J. B., Fox, P. T., & Eickhoff, S. B. (2012). Modelling neural correlates of working memory: a coordinate-based meta-analysis. NeuroImage, 60(1), 830–846. 10.1016/j.neuroimage.2011.11.050

[15] Jung, K., Min, Y., & Han, S. W. (2021). Response of multiple demand network to visual search demands. NeuroImage, 229, 117755. 10.1016/j.neuroimage.2021.117755

[16] Moerel, D., Rich, A. N., & Woolgar, A. (2024). Selective attention and decision-making have separable neural bases in space and time. Journal of Neuroscience, 44(38), 1–13. Article e0224242024. 10.1523/JNEUROSCI.0224-24.2024

[17] Niendam, T. A., Laird, A. R., Ray, K. L., Dean, Y. M., Glahn, D. C., & Carter, C. S. (2012). Meta-analytic evidence for a superordinate cognitive control network subserving diverse executive functions. Cognitive, Affective, & Behavioral Neuroscience, 12(2), 241–268. 10.3758/s13415-011-0083-5

[18] Gonzalez Alam, T., Murphy, C., Smallwood, J., & Jefferies, E. (2018). Meaningful inhibition: Exploring the role of meaning and modality in response inhibition. NeuroImage, 181, 108–119. 10.1016/j.neuroimage.2018.06.074

[19] Vincent, J. L., Kahn, I., Snyder, A. Z., Raichle, M. E., & Buckner, R. L. (2008). Evidence for a frontoparietal control system revealed by intrinsic functional connectivity. Journal of Neurophysiology, 100(6), 3328–3342. 10.1152/jn.90355.2008

[20] Yeo, B. T., Krienen, F. M., Sepulcre, J., Sabuncu, M. R., Lashkari, D., Hollinshead, M., Roffman, J. L., Smoller, J. W., Zöllei, L., Polimeni, J. R., Fischl, B., Liu, H., & Buckner, R. L. (2011). The organization of the human cerebral cortex estimated by intrinsic functional connectivity. Journal of Neurophysiology, 106(3), 1125–1165. 10.1152/jn.00338.2011

[21] Power, J. D., Cohen, A. L., Nelson, S. M., Wig, G. S., Barnes, K. A., Church, J. A., Vogel, A. C., Laumann, T. O., Miezin, F. M., Schlaggar, B. L., & Petersen, S. E. (2011). Functional network organization of the human brain. Neuron, 72(4), 665–678. 10.1016/j.neuron.2011.09.006

[22] Ji, J. L., Spronk, M., Kulkarni, K., Repov̌s, G., Anticevic, A., & Cole, M. W. (2019). Mapping the human brain’s cortical-subcortical functional network organization. NeuroImage, 185, 35–57. 10.1016/j.neuroimage.2018.10.006

[23] Assem, M., Glasser, M. F., Van Essen, D. C., & Duncan, J. (2020). A domain-general cognitive core defined in multimodally parcellated human cortex. Cerebral Cortex, 30(8), 4361–4380. 10.1093/cercor/bhaa040

[24] Cole, M. W., & Schneider, W. (2007). The cognitive control network: Integrated cortical regions with dissociable functions. NeuroImage, 37(1), 343–360. 10.1016/j.neuroimage.2007.04.046

[25] Marek, S., & Dosenbach, N. U. F. (2018). The frontoparietal network: Function, electrophysiology, and importance of individual precision mapping. Dialogues in Clinical Neuroscience, 20(2), 133–140. 10.31887/DCNS.2018.20.2/smarek

[26] Hugdahl, K., Raichle, M. E., Mitra, A., & Specht, K. (2015). On the existence of a generalized non-specific task-dependent network. Frontiers in Human Neuroscience, 9, 430. 10.3389/fnhum.2015.00430

[27] Lu, R., Dermody, N., Duncan, J., & Woolgar, A. (2024). Aperiodic and oscillatory systems underpinning human domain-general cognition. Communications Biology, 7, 1643. 10.1038/s42003-024-07397-7

[28] Lu, R. (2025). Linking the multiple-demand cognitive control system to human electrophysiological activity. Neuropsychologia, 210, 109096. 10.1016/j.neuropsychologia.2025.109096

[29] Hartman, S., Arnon, T., & Erez, Y. (2025). Individual-level neuroimaging of cognitive control: From basic science to brain tumor clinical applications. Neuropsychologia, 217, 109207. 10.1016/j.neuropsychologia.2025.109207

[30] Duncan, J., Burgess, P., & Emslie, H. (1995). Fluid intelligence after frontal lobe lesions. Neuropsychologia, 33(3), 261–268. 10.1016/0028-3932(94)00124-8

[31] Woolgar, A., Parr, A., Cusack, R., Thompson, R., Nimmo-Smith, I., Torralva, T., … & Duncan, J. (2010). Fluid intelligence loss linked to restricted regions of damage within frontal and parietal cortex. Proceedings of the National Academy of Sciences, 107(33), 14899–14902. 10.1073/pnas.1007928107

[32] Woolgar, A., Duncan, J., Manes, F., & Fedorenko, E. (2018). Fluid intelligence is supported by the multiple-demand system, not the language system. Nature Human Behaviour, 2(3), 200–204. 10.1038/s41562-017-0282-3.

[33] Gläscher, J., Tranel, D., Paul, L. K., Rudrauf, D., Rorden, C., Hornaday, A., … & Adolphs, R. (2009). Lesion mapping of cognitive abilities linked to intelligence. Neuron, 61(5), 681–691. 10.1016/j.neuron.2009.01.026

[34] Roca, M., Parr, A., Thompson, R., Woolgar, A., Torralva, T., Antoun, N., … & Duncan, J. (2010). Executive function and fluid intelligence after frontal lobe lesions. Brain, 133(1), 234–247. 10.1093/brain/awp269

[35] Siegel, J. S., Ramsey, L. E., Snyder, A. Z., Metcalf, N. V., Chacko, R. V., Weinberger, K., Baldassarre, A., Hacker, C. D., Shulman, G. L., & Corbetta, M. (2016). Disruptions of network connectivity predict impairment in multiple behavioral domains after stroke. Proceedings of the National Academy of Sciences of the United States of America, 113(30), E4367–E4376. 10.1073/pnas.1521083113

[36] Woolgar, A., Bor, D., & Duncan, J. (2013). Global increase in task-related fronto-parietal activity after focal frontal lobe lesion. Journal of Cognitive Neuroscience, 25(9), 1542–1552. 10.1162/jocn_a_00408

[37] Cope, T. E., Hughes, L. E., Phillips, H. N., Adams, N. E., Jafarian, A., Nesbitt, D., Assem, M., Woolgar, A., Duncan, J., & Rowe, J. B. (2022). Causal Evidence for the Multiple Demand Network in Change Detection: Auditory Mismatch Magnetoencephalography across Focal Neurodegenerative Diseases. The Journal of neuroscience : the official journal of the Society for Neuroscience, 42(15), 3197–3215. 10.1523/JNEUROSCI.1622-21.2022

[38] Woolgar, A., Hampshire, A., Thompson, R., & Duncan, J. (2011). Adaptive coding of task-relevant information in human frontoparietal cortex. Journal of Neuroscience, 31(41), 14592–14599. 10.1523/JNEUROSCI.3524-10.2011

[39] Woolgar, A., Jackson, J., & Duncan, J. (2016). Coding of visual, auditory, rule, and response information in the brain: 10 years of multivoxel pattern analysis. Journal of Cognitive Neuroscience, 28(10), 1433–1454. 10.1162/jocn_a_00964

[40] Woolgar, A., & Zopf, R. (2017). Multisensory coding in the multiple-demand regions: Vibro-tactile task information is coded in frontoparietal cortex. Journal of Neurophysiology, 118(2), 703–716. 10.1152/jn.00559.2016

[41] Crittenden, B. M., Mitchell, D. J., & Duncan, J. (2016). Task encoding across the multiple demand cortex is consistent with a frontoparietal and cingulo-opercular dual networks distinction. Journal of Neuroscience, 36(23), 6147–6155. 10.1523/JNEUROSCI.4590-15.2016

[42] Pischedda, D., Görgen, K., Haynes, J.-D., & Reverberi, C. (2017). Neural representations of hierarchical rule sets: The human control system represents rules irrespective of the hierarchical level. Journal of Neuroscience, 37(50), 12281–12296. 10.1523/JNEUROSCI.3088-16.2017

[43] Schultz, D. H., Ito, T., & Cole, M. W. (2022). Global connectivity fingerprints predict the domain generality of multiple-demand regions. Cerebral Cortex, 32(20), 4464–4479. 10.1093/cercor/bhab495

[44] Vaidya, A. R., Jones, H. M., Castillo, J., & Badre, D. (2021). Neural representation of abstract task structure during generalization. eLife, 10, e63226. 10.7554/eLife.63226

[45] Jackson, J. B., & Woolgar, A. (2018). Adaptive coding in the human brain: Distinct object features are encoded by overlapping voxels in frontoparietal cortex. Cortex; a journal devoted to the study of the nervous system and behavior, 108, 25–34. 10.1016/j.cortex.2018.07.006

[46] Cocuzza, C. V., Ito, T., Schultz, D., Bassett, D. S., & Cole, M. W. (2020). Flexible Coordinator and Switcher Hubs for Adaptive Task Control. The Journal of neuroscience : the official journal of the Society for Neuroscience, 40(36), 6949–6968. 10.1523/JNEUROSCI.2559-19.2020

[47] Nee, D. E., Brown, J. W., Askren, M. K., Berman, M. G., Demiralp, E., Krawitz, A., & Jonides, J. (2013). A meta-analysis of executive components of working memory. Cerebral cortex (New York, N.Y. : 1991), 23(2), 264–282. 10.1093/cercor/bhs007

[48] Rodŕıguez-Nieto, G., Seer, C., Sidlauskaite, J., Vleugels, L., Van Roy, A., Hardwick, R. M., & Swinnen, S. P. (2022). Inhibition, shifting and updating: Inter- and intra-domain commonalities and differences from an executive functions activation likelihood estimation meta-analysis. NeuroImage, 264, 119665. 10.1016/j.neuroimage.2022.119665

[49] Numssen, O., Bzdok, D., & Hartwigsen, G. (2021). Functional specialization within the inferior parietal lobes across cognitive domains. eLife, 10, e63591. 10.7554/eLife.63591

[50] Assem, M., Shashidhara, S., Glasser, M. F., & Duncan, J. (2024). Basis of executive functions in fine-grained architecture of cortical and subcortical human brain networks. Cerebral Cortex, 34(2), bhad537. 10.1093/cercor/bhad537

[51] Braga, R. M., Leech, R., & Sharp, D. J. (2017). Parallel interdigitated distributed networks within the individual estimated by intrinsic functional connectivity. Neuron, 95(2), 457–471.e5. 10.1016/j.neuron.2017.06.027

[52] Murphy, A.C., Bertolero, M.A., Papadopoulos, L. et al. Multimodal network dynamics underpinning working memory. Nat Commun 11, 3035 (2020). 10.1038/s41467-020-15541-0

[53] Dixon, M. L., De La Vega, A., Mills, C., Andrews-Hanna, J., Spreng, R. N., Cole, M. W., & Christoff, K. (2018). Heterogeneity within the frontoparietal control network and its relationship to the default and dorsal attention networks. Proceedings of the National Academy of Sciences, 115(13), E3068–E3077. 10.1073/pnas.1715766115

[54] Beaty, R. E., et al. (2021). Functional realignment of frontoparietal subnetworks during creative cognition. Cerebral Cortex, 31(10), 4464–4477. 10.1093/cercor/bhab055

[55] Camilleri, J. A., et al. (2018). Definition and characterization of an extended multiple-demand network (eMDN). NeuroImage, 165, 138–147. 10.1016/j.neuroimage.2017.10.032

[56] Glasser, M. F., Coalson, T. S., Robinson, E. C., Hacker, C. D., Harwell, J., Yacoub, E., Ugurbil, K., Andersson, J., Beckmann, C. F., Jenkinson, M., Smith, S. M., & Van Essen, D. C. (2016). A multi-modal parcellation of human cerebral cortex. Nature, 536(7615), 171–178. 10.1038/nature18933

[57] Satopää, V. A., Albrecht, J., Irwin, D., Raghavan, B., & Irwin, D. (2011). Finding a “kneedle” in a haystack: Detecting knee points in system behavior. Proceedings of the 31st International Conference on Distributed Computing Systems Workshops, 166–171. 10.1109/ICDCSW.2011.20

[58] Finn, E. S., Bandettini, P. A., Shen, X., & Constable, R. T. (2021). Movie-watching outperforms rest for functional connectivity–based prediction of behavior. NeuroImage, 235, 118018. 10.1016/j.neuroimage.2021.118018

[59] Meer, J. N. van der, Breakspear, M., Chang, L. J., & Sonkusare, S. (2020). Movie viewing elicits rich and reliable brain state dynamics. Nature Communications, 11(1), 5000. 10.1038/s41467-020-18717-w

[60] Vanderwal, T., Eilbott, J., & Castellanos, F. X. (2019). Movies in the magnet: Naturalistic paradigms in developmental functional neuroimaging. Developmental cognitive neuroscience, 36, 100600. 10.1016/j.dcn.2018.10.004

[61] Rajimehr, R., Xu, H., Farahani, A., Kornblith, S., Duncan, J., & Desimone, R. (2024). Functional architecture of cerebral cortex during naturalistic movie watching. Neuron, 112(24), 4130–4146.e3. 10.1016/j.neuron.2024.10.005

[62] “What” and “where” in the human brain. Current Opinion in Neurobiology, 4(2), 157–165. 10.1016/0959-4388(94)90066-3

[63] Separate visual pathways for perception and action. Trends in Neurosciences, 15(1), 20–25. 10.1016/0166-2236(92)90344-8

[64] Rolls, E. T. (2024). Two what, two where, visual cortical streams in humans. Neuroscience & Biobehavioral Reviews, 160, Article 105650. 10.1016/j.neubiorev.2024.105650

[65] Corbetta, M., & Shulman, G. L. (2002). Control of goal-directed and stimulus-driven attention in the brain. Nature Reviews Neuroscience, 3(3), 201–215. 10.1038/nrn755

[66] Fox, M. D., Corbetta, M., Snyder, A. Z., Vincent, J. L., & Raichle, M. E. (2006). Spontaneous neuronal activity distinguishes human dorsal and ventral attention systems. Proceedings of the National Academy of Sciences, 103(26), 10046–10051. 10.1073/pnas.0604187103

[67] Fruchterman, T. M. J., & Reingold, E. M. (1991). Graph drawing by force-directed placement. Software: Practice and Experience, 21(11), 1129–1164. 10.1002/spe.4380211102

[68] Freeman, L. C. (1977). A set of measures of centrality based on betweenness. Sociometry, 40(1), 35–41. 10.2307/3033543

[69] Leydesdorff, L. (2007). Betweenness centrality as an indicator of the interdisciplinarity of scientific journals. Journal of the American Society for Information Science and Technology, 58(9), 1303–1319. 10.1002/asi.20614

[70] Hwang, W. C., Zhang, A., & Ramanathan, M. (2008). Identification of information flow-modulating drug targets: a novel bridging paradigm for drug discovery. Clinical pharmacology and therapeutics, 84(5), 563–572. 10.1038/clpt.2008.129

[71] Meghanathan, Natarajan. Neighborhood-based bridge node centrality tuple for complex network analysis. Applied Network Science, 6 (1). Retrieved from https://par.nsf.gov/biblio/10319137. 10.1007/s41109-021-00388-1

[72] Rajeh, S., Savonnet, M., Leclercq, E., & Cherifi, H. (2021). Characterizing the interactions between classical and community-aware centrality measures in complex networks. Scientific reports, 11(1), 10088. 10.1038/s41598-021-89549-x

[73] Steyvers, M., & Schafer, R. J. (2020). Inferring latent learning factors in large-scale cognitive training data. Nature human behaviour, 4(11), 1145–1155. 10.1038/s41562-020-00935-3

[74] Tipping, M. E., & Bishop, C. M. (1999). Probabilistic principal component analysis. Journal of the Royal Statistical Society: Series B (Statistical Methodology), 61(3), 611–622. 10.1111/1467-9868.00196

[75] Ilin, A., & Raiko, T. (2010). Practical approaches to principal component analysis in the presence of missing values. Journal of Machine Learning Research, 11, 1957–2000. http://jmlr.org/papers/v11/ilin10a.html

[76] Kovacs, K., & Conway, A. R. A. (2016). Process overlap theory: A unified account of the general factor of intelligence. Psychological Inquiry, 27(3), 151–177. 10.1080/1047840X.2016.1153946

[77] Osman, A. M., Jaffe, P. I., Ng, N. F., Kerlan, K. R., & Schafer, R. J. (2023). Transfer of learning: Analysis of dose-response functions from a large-scale, online, cognitive training dataset. PloS one, 18(5), e0281095. 10.1371/journal.pone.0281095

[78] Zhang, Z., Peng, P., Eickhoff, S. B., Lin, X., Zhang, D., & Wang, Y. (2021). Neural substrates of the executive function construct, age-related changes, and task materials in adolescents and adults: ALE meta-analyses of 408 fMRI studies. Developmental science, 24(6), e13111. 10.1111/desc.13111

[79] Witt, S. T., van Ettinger-Veenstra, H., Salo, T., Riedel, M. C., & Laird, A. R. (2021). What Executive Function Network is that? An Image-Based Meta-Analysis of Network Labels. Brain topography, 34(5), 598–607. 10.1007/s10548-021-00847-z

[80] Power, J. D., Schlaggar, B. L., Lessov-Schlaggar, C. N., & Petersen, S. E. (2013). Evidence for hubs in human functional brain networks. Neuron, 79(4), 798–813. 10.1016/j.neuron.2013.07.035

[81] Bertolero, M. A., Yeo, B. T. T., & D’Esposito, M. (2015). The modular and integrative functional architecture of the human brain. Proceedings of the National Academy of Sciences of the United States of America, 112(49), E6798–E6807. 10.1073/pnas.1510619112

[82] Margulies, D. S., Ghosh, S. S., Goulas, A., Falkiewicz, M., Huntenburg, J. M., Langs, G., Bezgin, G., Eickhoff, S. B., Castellanos, F. X., Petrides, M., Jefferies, E., & Smallwood, J. (2016). Situating the default-mode network along a principal gradient of macroscale cortical organization. Proceedings of the National Academy of Sciences of the United States of America, 113(44), 12574–12579. 10.1073/pnas.1608282113

[83] Huntenburg, J. M., Bazin, P. L., & Margulies, D. S. (2018). Large-Scale Gradients in Human Cortical Organization. Trends in cognitive sciences, 22(1), 21–31. 10.1016/j.tics.2017.11.002

[84] Choi, E. Y., Drayna, G. K., & Badre, D. (2018). Evidence for a Functional Hierarchy of Association Networks. Journal of cognitive neuroscience, 30(5), 722–736. 10.1162/jocn_a_01229

[85] Nee, D. E., & D’Esposito, M. (2016). The hierarchical organization of the lateral prefrontal cortex. eLife, 5, e12112. 10.7554/eLife.12112

[86] Nee, D. E., & D’Esposito, M. (2017). Causal evidence for lateral prefrontal cortex dynamics supporting cognitive control. eLife, 6, e28040. 10.7554/eLife.28040

[87] Nee D. E. (2021). Integrative frontal-parietal dynamics supporting cognitive control. eLife, 10, e57244. 10.7554/eLife.57244

[88] Chiou, R., Jefferies, E., Duncan, J., Humphreys, G. F., & Lambon Ralph, M. A. (2023). A middle ground where executive control meets semantics: The neural substrates of semantic control are topographically sandwiched between the multiple-demand and default-mode systems. Cerebral Cortex, 33(8), 4512–4526. 10.1093/cercor/bhac358

[89] Binder, J. R., Desai, R. H., Graves, W. W., & Conant, L. L. (2009). Where is the semantic system? A critical review and meta-analysis of 120 functional neuroimaging studies. Cerebral Cortex, 19(12), 2767–2796. 10.1093/cercor/bhp055

[90] Baldassano, C., Fei-Fei, L., & Beck, D. M. (2016). Two distinct scene-processing networks connecting vision and memory. Cerebral Cortex, 26(7), 2815–2828. 10.1093/cercor/bhv165

[91] A posterior–anterior distinction between scene perception and spatial memory in the human medial parietal cortex. Journal of Neuroscience, 39(14), 2663–2677. 10.1523/JNEUROSCI.1939-18.2019

[92] Collins, J. A., & Olson, I. R. (2014). Beyond the FFA: The role of the ventral anterior temporal lobes in face processing. Neuropsychologia, 61, 65–79. 10.1016/j.neuropsychologia.2014.06.005

[93] Popham, S. F., Huth, A. G., Bilenko, N. Y., Deniz, F., Gao, J. S., Nunez-Elizalde, A. O., & Gallant, J. L. (2021). Visual and linguistic semantic representations are aligned at the border of human visual cortex. Nature neuroscience, 24(11), 1628–1636. 10.1038/s41593-021-00921-6

[94] Spearman, C. (1904). “General intelligence,” objectively determined and measured. The American Journal of Psychology, 15(2), 201–292. 10.2307/1412107

[95] Carroll, J. B. (1993). Human cognitive abilities: A survey of factor-analytic studies. Cambridge University Press. 10.1017/CBO9780511571312

[96] Hampshire, A., Highfield, R. R., Parkin, B. L., & Owen, A. M. (2012). Fractionating human intelligence. Neuron, 76(6), 1225–1237. 10.1016/j.neuron.2012.06.022

[97] Sripada, C., Angstadt, M., Rutherford, S., Taxali, A., & Shedden, K. (2020). Toward a “treadmill test” for cognition: Improved prediction of general cognitive ability from the task-activated brain. Human Brain Mapping. Advance online publication. 10.1002/hbm.25007

[98] Höffler, T. N., Kóc-Januchta, M., & Leutner, D. (2017). More Evidence for Three Types of Cognitive Style: Validating the Object-Spatial Imagery and Verbal Questionnaire Using Eye Tracking when Learning with Texts and Pictures. Applied cognitive psychology, 31(1), 109–115. 10.1002/acp.3300

[99] Cattell, R. B. (1963). Theory of fluid and crystallized intelligence: A critical experiment. Journal of Educational Psychology, 54(1), 1–22. 10.1037/h0046743

[100] Horn, J. L., & Cattell, R. B. (1967). Age differences in fluid and crystallized intelligence. Acta Psychologica, 26, 107–129. 10.1016/0001-6918(67)90011-X

[101] Jung, R. E., & Haier, R. J. (2007). The parieto-frontal integration theory (P-FIT) of intelligence: Converging neuroimaging evidence. Behavioral and Brain Sciences, 30(2), 135–154. 10.1017/S0140525X07001185

[102] Kim, G., Kim, DK. & Jeong, H. Spontaneous emergence of rudimentary music detectors in deep neural networks. Nat Commun 15, 148 (2024). 10.1038/s41467-023-44516-0

[103] Van Essen, D. C., Ugurbil, K., Auerbach, E., Barch, D., Behrens, T. E., Bucholz, R., Chang, A., Chen, L., Corbetta, M., Curtiss, S. W., Della Penna, S., Feinberg, D., Glasser, M. F., Harel, N., Heath, A. C., Larson-Prior, L., Marcus, D., Michalareas, G., Moeller, S., Oostenveld, R., . . . WU-Minn HCP Consortium (2012). The Human Connectome Project: a data acquisition perspective. NeuroImage, 62(4), 2222–2231. 10.1016/j.neuroimage.2012.02.018

[104] Glasser, M. F., Sotiropoulos, S. N., Wilson, J. A., Coalson, T. S., Fischl, B., Andersson, J. L., Xu, J., Jbabdi, S., Webster, M., Polimeni, J. R., Van Essen, D. C., Jenkinson, M., & WU-Minn HCP Consortium (2013). The minimal preprocessing pipelines for the Human Connectome Project. NeuroImage, 80, 105–124. 10.1016/j.neuroimage.2013.04.127

[105] Jenkinson, M., Beckmann, C. F., Behrens, T. E., Woolrich, M. W., & Smith, S. M. (2012). FSL. NeuroImage, 62(2), 782–790. 10.1016/j.neuroimage.2011.09.015

[106] Salimi-Khorshidi, G., Douaud, G., Beckmann, C. F., Glasser, M. F., Griffanti, L., & Smith, S. M. (2014). Automatic denoising of functional MRI data: combining independent component analysis and hierarchical fusion of classifiers. NeuroImage, 90, 449–468. 10.1016/j.neuroimage.2013.11.046

[107] Griffanti, L., Salimi-Khorshidi, G., Beckmann, C. F., Auerbach, E. J., Douaud, G., Sexton, C. E., Zsoldos, E., Ebmeier, K. P., Filippini, N., Mackay, C. E., Moeller, S., Xu, J., Yacoub, E., Baselli, G., Ugurbil, K., Miller, K. L., & Smith, S. M. (2014). ICA-based artefact removal and accelerated fMRI acquisition for improved resting state network imaging. NeuroImage, 95, 232–247. 10.1016/j.neuroimage.2014.03.034

[108] Fischl B. (2012). FreeSurfer. NeuroImage, 62(2), 774–781. 10.1016/j.neuroimage.2012.01.021

[109] Marcus, D. S., Harwell, J., Olsen, T., Hodge, M., Glasser, M. F., Prior, F., Jenkinson, M., Laumann, T., Curtiss, S. W., & Van Essen, D. C. (2011). Informatics and data mining tools and strategies for the human connectome project. Frontiers in neuroinformatics, 5, 4. 10.3389/fninf.2011.00004

[110] Glasser, M. F., Goyal, M. S., Preuss, T. M., Raichle, M. E., & Van Essen, D. C. (2014). Trends and properties of human cerebral cortex: correlations with cortical myelin content. NeuroImage, 93 Pt 2, 165–175. 10.1016/j.neuroimage.2013.03.060

[111] Glasser, M. F., & Van Essen, D. C. (2011). Mapping human cortical areas in vivo based on myelin content as revealed by T1- and T2-weighted MRI. The Journal of neuroscience : the official journal of the Society for Neuroscience, 31(32), 11597–11616. 10.1523/JNEUROSCI.2180-11.2011

[112] Robinson, E. C., Garcia, K., Glasser, M. F., Chen, Z., Coalson, T. S., Makropoulos, A., Bozek, J., Wright, R., Schuh, A., Webster, M., Hutter, J., Price, A., Cordero Grande, L., Hughes, E., Tusor, N., Bayly, P. V., Van Essen, D. C., Smith, S. M., Edwards, A. D., Hajnal, J., . . . Rueckert, D. (2018). Multimodal surface matching with higher-order smoothness constraints. NeuroImage, 167, 453–465. 10.1016/j.neuroimage.2017.10.037

